# Mutant p53 suppresses innate immune signaling to promote tumorigenesis

**DOI:** 10.1101/2020.03.12.989384

**Authors:** Monisankar Ghosh, Suchandrima Saha, Julie Bettke, Rachana Nagar, Alejandro Parrales, Tomoo Iwakuma, Adrianus W. M. van der Velden, Luis A. Martinez

## Abstract

Mutations in the p53 tumor suppressor occur very frequently in human cancer. Often, such mutations lead to the constitutive overproduction of mutant p53 (mtp53) proteins, which can exert a cancer-promoting gain-of-function (GOF). We have identified a novel mechanism by which mtp53 controls both cell-autonomous and non-cell autonomous signaling to promote cancer cell survival and suppress tumor immune surveillance. Mtp53 interferes with the function of the cytoplasmic DNA sensing machinery, cGAS-STING-TBK1-IRF3, that controls the activation of the innate immune response. We find that mtp53, but not wildtype p53, binds to TANK binding protein kinase (TBK1) and inhibits both its basal and agonist-induced activity. The association of mtp53 with TBK1 prevents the formation of a trimeric complex between TBK1-STING-IRF3, which is required for activation, nuclear translocation and transcriptional activity of IRF3. Mtp53 knockdown restores TBK1 activity, resulting in the transcriptional induction of IRF3 target genes and IRF3-dependent apoptosis. Furthermore, inactivation of innate immune signaling by mtp53 alters cytokine production resulting in immune evasion. Restored TBK1 signaling was sufficient to bypass mtp53 and reactivate cell-autonomous and non-cell autonomous tumor control. Thus, overriding mtp53’s inhibition of this cytosolic DNA sensing pathway may ultimately lead to restored immune cell function and cancer cell eradication.

## Introduction

The tumor suppressor p53, is one of the most important proteins that protect the genome from sustaining DNA damage and propagating genetic lesions into daughter cells. It has long been recognized that wildtype p53 functions as the guardian of the genome because it promotes genomic stability through various pathways and its loss results in genomic instability.(*1, 2*) In addition, wildtype p53 can induce a senescent program that promotes the formation of a tumor suppressive microenvironment in a non-cell autonomous manner(*3*). Moreover, loss of wildtype p53 can trigger the activation of the WNT pathway, resulting in systemic inflammation that drives cancer metastasis.(*4*). Given the high degree of genetic alterations in cancer, it is not surprising that TP53 is frequently inactivated.(*5*) The most common form of genetic lesions in the TP53 gene are missense point mutations in the DNA binding domain.(*5*) These missense mutations typically inactivate p53’s tumor suppressor activity while simultaneously generating an oncogenic mtp53 protein that exhibits gain-of function activities.(*5*) Recently, it has been reported that 91% of cancers that have a mtp53 allele have lost their second allele through mutation or chromosomal loss.(*6*) Furthermore, the presence of mtp53 correlates with increased chromosomal instability leading to loss of tumor suppressor genes and amplification of oncogenes.(*6, 7*)

Aneuploidy is considered one of the hallmarks of cancer and is thought to play an important role in driving tumor cell evolution.(*8*) Aneuploidy has been shown to provoke cell cycle arrest, senescence, increased pro-inflammatory cytokine production and upregulation of Natural Killer (NK) cell ligands in tumor cells, resulting in their subsequent killing by NK cells.(*9*) In an *in vivo* setting, the increased production of pro-inflammatory cytokines coupled with increased NK cell ligand expression permits the recruitment of immune cells and clearance of abnormal cells.(*10*) Thus, in addition to the cell-autonomous mechanisms that suppress genetic instability, cells also utilize non-cell autonomous mechanisms to signal their own destruction.(*9*) However, chromosomal instability can give rise to micronuclei, which are prone to rupturing and releasing the DNA into the cytoplasm.(*11*) DNA leaked into the cytoplasm is recognized by the DNA binding protein, cyclic GMP-AMP synthase (cGAS), which in turn triggers innate immune signaling that results in the production of type I interferons.(*12*) Mechanistically, DNA promotes cGAS homodimerization and production of the second messenger cyclic-GMP-AMP (cGAMP).(*12, 13*) This cGAMP molecule is then recognized by the endoplasmic resident, Stimulator of Interferon Genes (STING), which then interacts with TANK binding kinase (TBK1).(*13, 14*) TBK1 auto-phosphorylates itself and phosphorylates STING to create a binding site for recruitment of the transcription factor, IRF3.(*14*) IRF3 forms a trimeric complex with STING and TBK1 and is then phosphorylated by TBK1 allowing it to homodimerize and translocate to the nucleus to regulate gene expression.(*13*) In addition to its function as a transcription factor, IRF3 can translocate to the mitochondria and promote pore formation by interacting with Bcl-xl.(*15*) Thus, the formation of the trimeric STING/TBK1/IRF3 complex is a prerequisite for TBK1 activation and downstream signaling by IRF3. The cGAS/STING/TBK1/IRF3 innate immune signaling pathway plays a key role in the suppression of tumor development through both cell autonomous and non-cell autonomous signaling resulting in immune cell-mediated tumor suppression.(*12, 16–18*) Cancer cells are known to have high levels of cytoplasmic DNA, which results in the constitutive (although still inducible) “basal” activation of the cGAS/STING pathway.(*19–21*) The cytosolic DNA comes from a variety of sources including ruptured micronuclei, reactivation of endogenous retroviral sequences, mitochondrial DNA, mitotic defects, etc..(*19–21*) In contrast, recent work has indicated that the cGAS/STING pathway can also promote metastasis in genetically unstable cells.(*20, 22*) Cytoplasmic DNA triggers the constitutive activation of cGAS and STING, however, in this setting IRF3 is not activated.(*20, 22*)

It remains unknown how signaling to IRF3 from cGAS/STING is disengaged despite the presence of cytoplasmic DNA in tumor cells.(*20*) It has previously been reported that there is a correlation between mtp53 and absence/reduced presence of immune cells in head and neck, and gastric cancers.(*23–25*) However, it is not known whether this is the functional outcome of wildtype p53 loss or a gain-of-function activity of mtp53. Since mtp53 is associated with genomic instability, we speculated that it may alter signaling through the cGAS/STING/TBK1/IRF3 pathway to permit the accumulation of cytoplasmic DNA without triggering IRF3 activation. We find that mtp53 binds to TBK1 and disrupts downstream signaling from cGAS/STING to TBK1, thereby preventing phosphorylation of its substrates. We demonstrate that in cells lacking p53, cytoplasmic DNA triggers IRF3 transcriptional activity and apoptosis. In contrast, mtp53 blunts the activation of IRF3 thereby promoting a tolerance for cytoplasmic DNA. Importantly, mtp53 promotes immune evasion by suppressing IRF3 activation *in vivo*. Thus, our work defines a novel gain-of-function activity of mtp53 by which it can block both cell autonomous and non-cell autonomous surveillance mechanisms thereby promoting cancer growth.

## Results

### Mtp53 spontaneously suppresses innate immune response

We sought to determine whether mtp53 regulates the innate immune signaling pathway, as an initial approach we performed shRNA knockdown of mtp53 in the breast cancer cell lines, BT549 (p53R249S) and MDA-MB-231 (p53R280K), and pancreatic cell lines, MiaPaca-2 (p53R248W) and KPC (p53R172H). In all 4 cell lines, we observed that mtp53 knockdown resulted in phosphorylation of TBK1 and its substrates, IRF3 and STING (Fig. 1A, 1B and Fig. S1A). Of note, the p53 targeting sequences are different for the mouse and human p53 and they both yielded a similar response. Furthermore, comparison of mouse embryonic fibroblasts (MEFs) derived from either p53 null (p53^-/-^) or mtp53 (p53^R172H/R172H^) expressing mice revealed that the presence of mtp53 correlated with decreased phosphorylation of TBK1, IRF3 and STING (Fig. 1C). We also engineered the 4T1 breast cancer cell line (p53 null) to express the R249S mtp53 and found that these cells had reduced phosphorylation of the TBK1 substrates (Fig. 1D). Next, we overexpressed mtp53 in two different, normal fibroblast cells, IMR-90 and human foreskin fibroblasts (HFF). In line with our observations, mtp53 (R280K) expression resulted in decreased phosphorylation of TBK1 and its substrates IRF3 and STING (Fig. S1B). These data suggest that mtp53 blocks the activity of innate immune signaling pathway.

**Figure 1:**
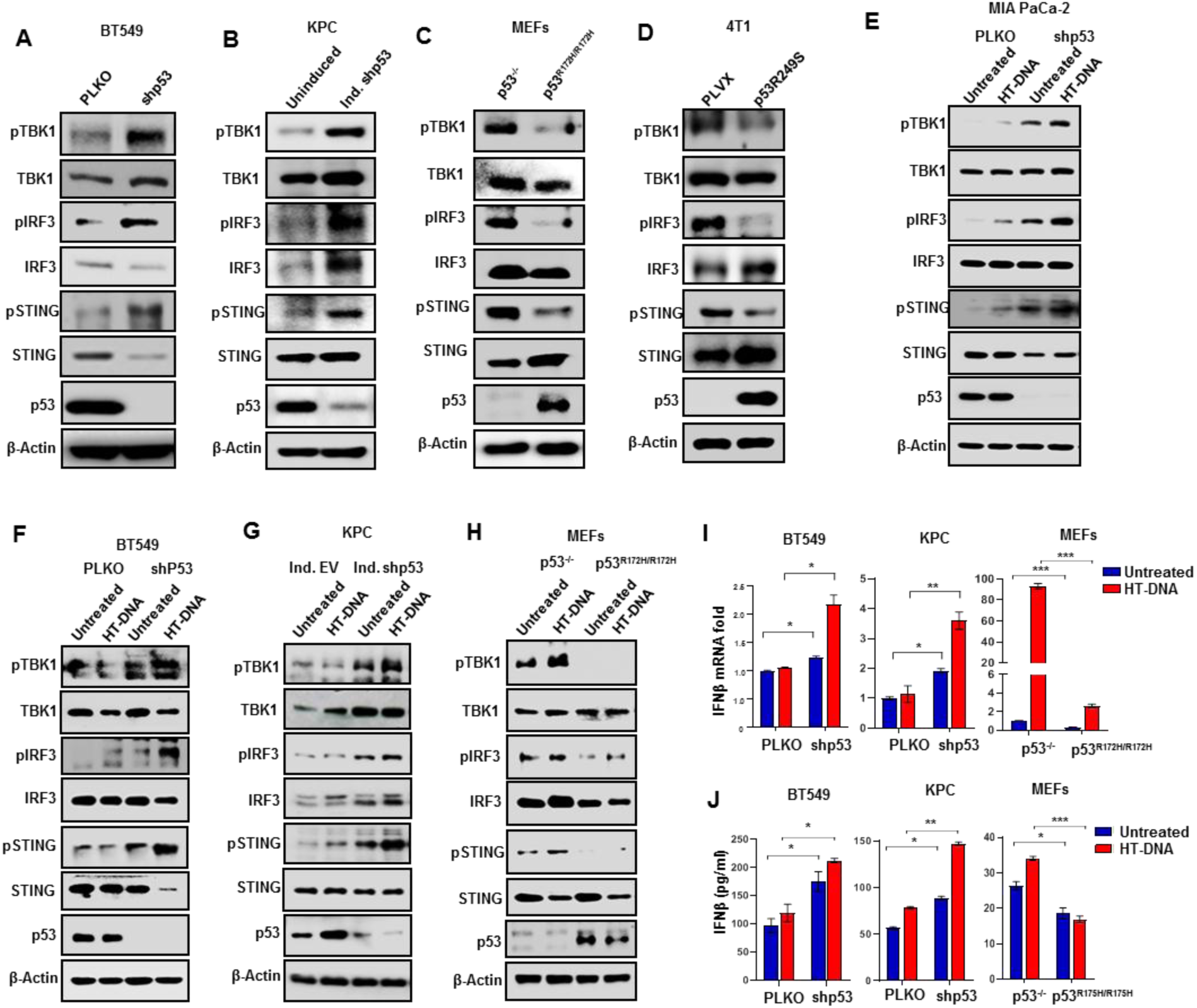
Mutant p53 spontaneously suppressed innate immune response. Immunoblot analysis of shRNA targeting mutant p53 in (A) BT549 and (B) KPC cells were subjected to western blotting analysis. (C) Immunoblot analysis of p53-/- and p53R172H/R172H MEFs. (D) p53 null 4T1 cells were engineered to express p53R249S and subjected to western blotting. (E, F and G) Indicated PLKO/ shp53 or induced shp53 cells were treated with 2 μg/ml of HT-DNA for 3 hrs, cells were harvested and subjected to western blot analysis. (H) p53-/- or p53R172H/R172H MEFs were treated with 2 μg/ml of HT-DNA for 3 hrs, cells were harvested and subjected to immunoblot analysis. (I and J) p53 knock down BT549 or KPC cells and mutant p53 expressing MEFs were treated with 2 μg/ml HT-DNA for 18 hrs, cells were harvested, RNA was isolated from the cells for RT-PCR and culture conditioned media was collected for ELISA to detect released IFNB1. Quantification graphs: In all panels, error bars represent mean with standard deviation. p values are based on Student’s t test. ∗∗∗p < 0.001, ∗∗p < 0.01, ∗p < 0.05, ns=non-significant.

Next, we wanted to ascertain if wildtype p53 could also regulate this pathway. Intriguingly, shRNA knockdown of wildtype p53 in A549 cells resulted in reduced phosphorylation of STING, TBK1, IRF3 (Fig. S1C). Moreover, in H1299 (p53 null) cells engineered to inducibly express wildtype p53, we observed the induction of TBK1, STING and IRF3 phosphorylation in response to p53 expression. (Fig. S1D) The phosphorylation of these proteins likely reflects the p53 dependent induction of IFI16, which cooperates with cGAS to activate TBK1/STING/IRF3.(*26–28*) (Supplementary Fig. S1D) In contrast, IFI16 levels are not affected by mtp53 expression (Fig. S1E). Thus, wildtype and mutant p53 function in an opposite manner in the control of the innate immune signaling pathway.

Activation of innate immune pathway triggers the type I interferon (IFNB1) response.(*29, 30*) Therefore, we tested whether mtp53-mediated changes in TBK1/STING/IRF3 phosphorylation correlated with activation of the IRF3 transcriptional target, IFNB1. We observed that mtp53 knockdown in BT549 and KPC induced IFNB1 mRNA, and conversely that ectopic mtp53 expression in 4T1 cells suppressed its expression (Fig. S1F). Furthermore, this phenotype was corroborated in p53^-/-^ MEFs which expressed higher levels of IFNB1 mRNA than their mtp53 (p53^R172H/R172H^) counterparts (Fig. S1F).

To determine if mtp53 also interferes with ligand-mediated activation of the STING-TBK1-IRF3 pathway, we transfected herring testis DNA (HT-DNA) in cells in which we reduced mtp53 levels by shRNA. Transfection of HT-DNA in BT549, MDA-MB-231, MIA PaCa-2, and KPC modestly activated the pathway. In contrast, shRNA knockdown of mutant p53 increased the basal phosphorylation levels of these proteins and these levels were further increased by HT-DNA treatment (Fig. 1E, 1F, 1G and Fig. S1G). In p53^-/-^ MEFs, we observed elevated levels of TBK1 substrate phosphorylation which could be moderately induced by HT-DNA. In contrast, phosphorylation of these substrates was barely detectable in the mtp53 MEFs, and was not induced by HT-DNA (Fig. 1H). Analysis of IFNB1 expression in response to HT-DNA revealed that BT549 and KPC cells failed to induce IFNB1 mRNA, whereas mtp53 knockdown resulted in robust induction in response to HT-DNA treatment (Fig. 1I). Consistent with the failure of mtp53 expressing MEFs to activate the pathway, these MEFs weakly induced IFNB1 expression in response to ligand, whereas the p53 null MEFs exhibited a strong induction (Fig. 1I). We also checked expression of various IRF3 controlled inflammatory cytokines in mtp53-knockdown BT549 and MDA-MB-231 cells treated with HT-DNA. Our results show that mtp53 knockdown augmented the induction of these cytokines in response to HT-DNA (Fig. S1H). All these cytokines are IRF3-target genes as IRF3 knockout in BT549 and MDA-MB-231 cells showed reduced mRNA of IFNB1, IFIT1, CXCL10, CCL5 and ISG15 (Fig. S2A and S2B). As anticipated, the reduced IFNB1 mRNA correlated with reduced IFNB1 protein secretion in the cultured medium as detected by ELISA. In BT549 and KPC cells, mtp53 knockdown resulted in increased IFNB1 secretion, whereas mtp53 expressing MEFs had reduced IFNB1 secretion (Fig. 1J).

To gain insight into what step of the pathway mtp53 was interfering with, we assessed if the STING agonist, cGAMP, could trigger IRF3 phosphorylation in mtp53 expressing cells. Transfection of cGAMP robustly induced IRF3 phosphorylation in the uninduced cells, but this did not occur in the cells induced to express mtp53 (Fig. S3A). These data suggested that mtp53 impedes signal transduction from STING to TBK1. TBK1 is activated by several pattern recognition receptor (PRR)-adaptor proteins that respond to viral RNA, cytosolic DNA and lipopolysaccharide (LPS), a component of the bacterial cell wall.(*31*) Intracellular double stranded RNA (dsRNA) is recognized by RIG-I, which activates the adaptor MAVS which subsequently activates TBK1.(*31*) Similarly, LPS is recognized by the toll like receptor 4 (TLR4), which activates TRIF, which in turn activates TBK1.(*31*) To assess if mtp53 only interfered with the response to cytoplasmic DNA, we treated cells with poly (I:C) (dsRNA) or LPS and assessed IRF3 phosphorylation. Treatment with either of these ligands induced IRF3 phosphorylation in cells not expressing mtp53, however, induction of mtp53 expression suppressed this response. (Supplementary Fig. S3A and S3B). Taken together, our data indicates that mtp53 blocks basal and PRR-induced TBK1 activation. Since these three PRR-adaptor proteins converge on TBK1 activation, this indicates that mtp53 blocks its activation.

### Mtp53 prevents nuclear localization of IRF3 and suppresses IRF3-dependent apoptosis

As mtp53 alters IRF3 activation and its targeted genes, next we investigated whether mtp53 affects IRF3 subcellular localization. IRF3 resides in the cytoplasm in unstimulated cells but upon activation of the STING-TBK1-IRF3 pathway, it dimerizes and translocate to the nucleus to induce its transcriptional targets. To assess how mtp53 regulates subcellular localization, we stably expressed GFP-IRF3 in H1299 cells that have a doxycycline inducible mtp53R248W. In uninduced cells, GFP-IRF3 was present in the cytoplasm and translocated to the nucleus after 3 hours of treatment with HT-DNA in approximately 50% of the cells. In contrast, mtp53 expression blunted the response to HT-DNA treatment, resulting in less than 20% of the cells containing nuclear GFP-IRF3 (Fig. 2A and 2B). Consistently, we found that whereas GFP-IRF3 remained mostly cytoplasmic in MDA-MB-231 cells infected with an empty vector (PLKO), shRNA knockdown of mtp53 resulted in a modest accumulation of GFP-IRF3 in the nucleus, and this was further increased by treatment with HT-DNA (Fig. S4A). We also observed that mutant p53 knockdown in BT549 and KPC cells spontaneously induced phospho-IRF3 accumulation in the nucleus (Fig. 2C)

**Figure 2:**
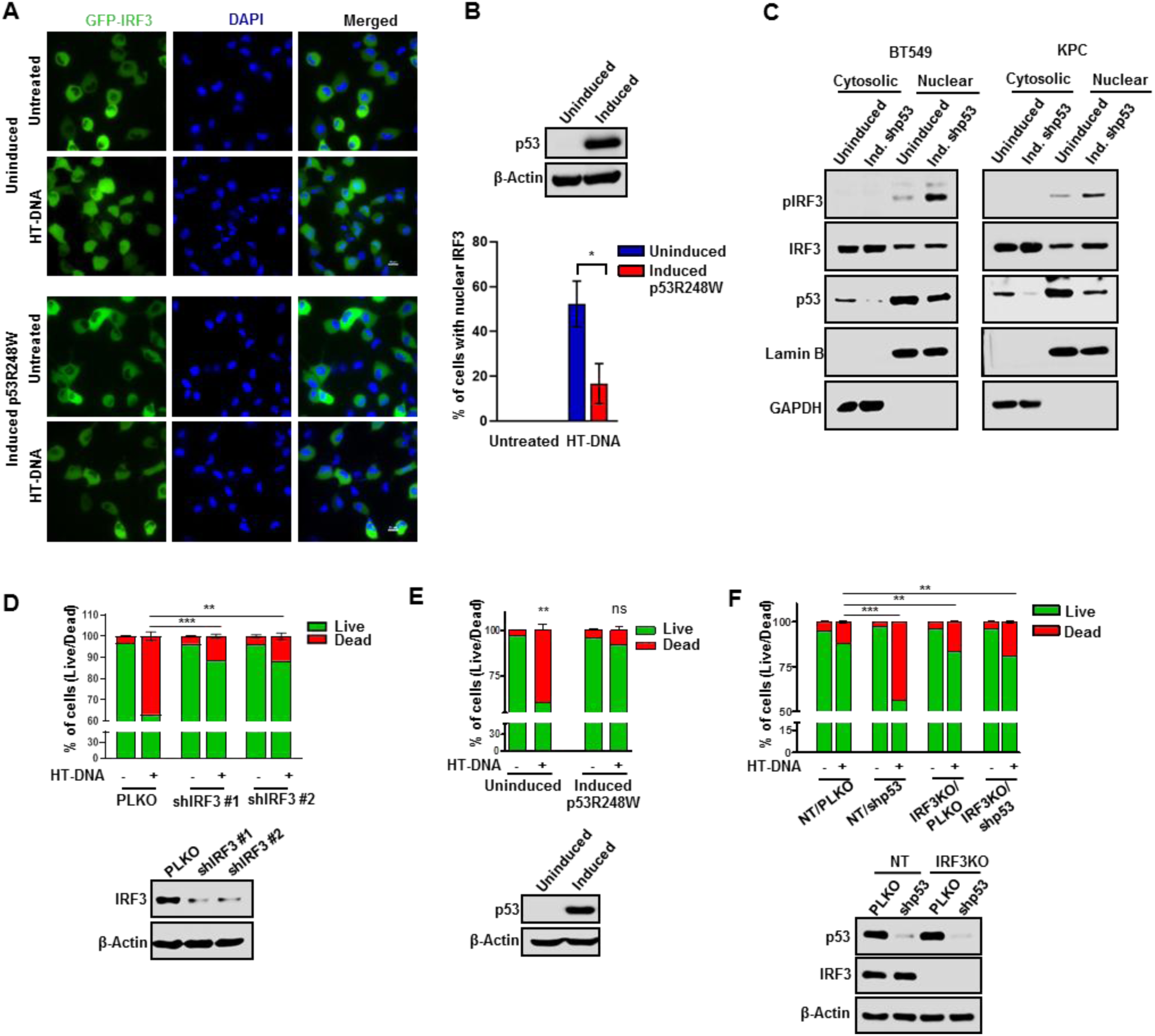
Mutant p53 blocks IRF3 nuclear translocation and suppresses IRF3 mediated apoptosis. (A) Representative fluorescent images of GFP-IRF3 positive H1299 cells were induced with Doxycycline for 24 hrs to express p53R248W. Cells were then transfected with 2 μg/ml of HT-DNA for 3 hrs and GFP-IRF3 was localization was visualized. Images were captured in Nikon Ti fluorescence microscopy at 60X magnification. (B) Immunoblots showing p53R248W expression. Quantification analysis of cells with nuclear IRF3 localization post HT-DNA treatment in uninduced and p53R248W induced H1299 cells. (C) Representative immunoblots of fractionated lysates of mutant p53 knock down BT549 and KPC cells. GAPDH and lamin B1 were used as loading controls for the cytoplasmic and nuclear fractions respectively. (D) Quantification analysis of apoptotic death in PLKO and shIRF3 H1299 cells treated with 2 μg/ml of HT-DNA for 24 hrs using flow cytometer. Immunoblots showing IRF3 knock down efficiency. (E) p53R248W induced H1299 cells were treated with 2 μg/ml of HT-DNA for 24 hrs. Cells were harvested, stained with Annexin V-FITC and PI and subjected to flow cytometry analysis. Graphical quantification showing that mutant p53 induction inhibited apoptotic death. Immunoblots showing p53R248W induction efficiency. (F) Quantitation of apoptosis in non-target (NT) and IRF3KO BT549 cells infected with either PLKO or shp53 virus, treated with 2 μg/ml HT-DNA for 24 hrs and apoptosis was analyzed using flow cytometry. Immunoblots showing p53 KD in NT and IRF3KO set. Quantification graphs: FoV= Field of View, (n=20). Scale bar = 20 um. In all panels, error bars represent mean with standard deviation. p values are based on Student’s t test. ∗∗∗p < 0.001, ∗∗p < 0.01, ns=non-significant.

Cytosolic DNA activates IRF3-dependent transcription, but can also induce mitochondria-mediated cell death.(*15, 17*) Specifically, IRF3 can interact with BAX and promote mitochondria pore formation and apoptosis.(*15*) In agreement, we observed that approximately 40% of H1299 cells treated with HT-DNA for 24 hours underwent apoptosis as determined by FACS analysis (Fig. 2D). This apoptotic response to HT-DNA was IRF3-dependent since it was largely abrogated by IRF3 knockdown with two different shRNAs (Fig. 2D). To further address if mtp53 antagonized this apoptotic response, we knocked down mtp53 in MDA-MB-231 cells and treated them with HT-DNA. FACS results showed that HT-DNA treatment resulted in apoptosis in about 30% of empty vector infected cells (PLKO), while the apoptotic response was more pronounced (approximately 70%) in the cells knocked down for mtp53 (Fig. S4B). Similarly, in uninduced H1299 cells, HT-DNA treatment resulted in apoptosis in approximately 40% of the population, and this was almost completely blocked by mtp53 expression (Fig. 2E). To provide further evidence that mtp53 could suppress HT-DNA induced IRF3-dependent apoptosis, we tested the response to HT-DNA of BT549 cells knocked down for mtp53 alone or in combination with IRF3 CRISPR knockout. Mtp53 knockdown increased the apoptotic response to HT-DNA from 10% to 60%, thereby supporting the notion that the mutant protein plays an important role in promoting cell survival in this context. Combining mtp53 knockdown and IRF3 knockout reduced the apoptotic response to ∼20%, indicating that the increased sensitivity of these cells to HT-DNA was largely IRF3-dependent (Fig. 2F). Collectively, these data indicate that mtp53-expressing cells fail to mount the cell intrinsic response to activation of the STING-TBK1-IRF3 pathway.

### Disruption of the TBK1/STING/IRF3 complex by mtp53 prevents IRF3 activation

IRF3 activation in response to cytoplasmic DNA requires TBK1 to phosphorylate STING, which provides a docking site for the recruitment of IRF3 and the formation of a trimeric complex that brings IRF3 in the proximity of TBK1, which then phosphorylates IRF3. To test if mtp53 blocks TBK1 phosphorylation of its substrates, we transfected all three (TBK1/STING/IRF3) with or without mtp53 and then analyzed their phosphorylation state. Ectopic TBK1 expression with STING and IRF3 in H1299 cells resulted in the phosphorylation of all three proteins, but induction of mtp53 resulted in their decreased phosphorylation (Fig. 3A). Our data suggested that mtp53 interferes with the function of TBK1 and thereby disrupts signaling to IRF3. We speculated that mtp53 could interact with one or more of these proteins, and thereby disrupt the formation of the complex required for IRF3 activation. Therefore, we tested the interaction of TBK1 with mutant and wildtype p53 in the H1299 inducible cells. Mtp53 interacted with TBK1, but wildtype p53 did not, despite its expression at equivalent levels (Fig. 3B). We also found that TBK1 interacted with endogenous mtp53 but not wildtype p53 (Fig. 3C). To determine how the mtp53 interaction with TBK1 impacts the formation of the trimeric complex, we co-transfected TBK1, STING and IRF3 in H1299 cells that were not induced or induced for mtp53 and then analyzed by immunoprecipitation/western blot the proteins that interacted with TBK1. Transfection of TBK1, STING and IRF3 in un-induced cells resulted in the co-precipitation of both STING and IRF3 with TBK1. Induction of mtp53 expression disrupted this interaction as indicated by their reduced co-immunoprecipitation with TBK1 (Fig. 3D). The active form of IRF3 functions as a dimer and thus we also determined if mtp53 impacted IRF3 homo-dimerization. To this end, we transfected GFP-IRF3 into the mtp53 inducible H1299 cells and immunoprecipitated with an anti-GFP antibody and did western blot with an anti-IRF3 antibody. In uninduced cells treated with HT-DNA, we detected the interaction of GFP-IRF3 with endogenous IRF3, indicating dimerization of IRF3, but induction of mtp53 disrupted this IRF3 complex (Fig. 3E). Of note, we did not detect an interaction between IRF3 and mtp53 suggesting that the IRF3 dimer is not directly disrupted by mtp53. Our data indicates that mtp53 prevents the formation of the trimeric TBK1/STING/IRF3 complex and thus precludes TBK1 and IRF3 activation.

**Figure 3:**
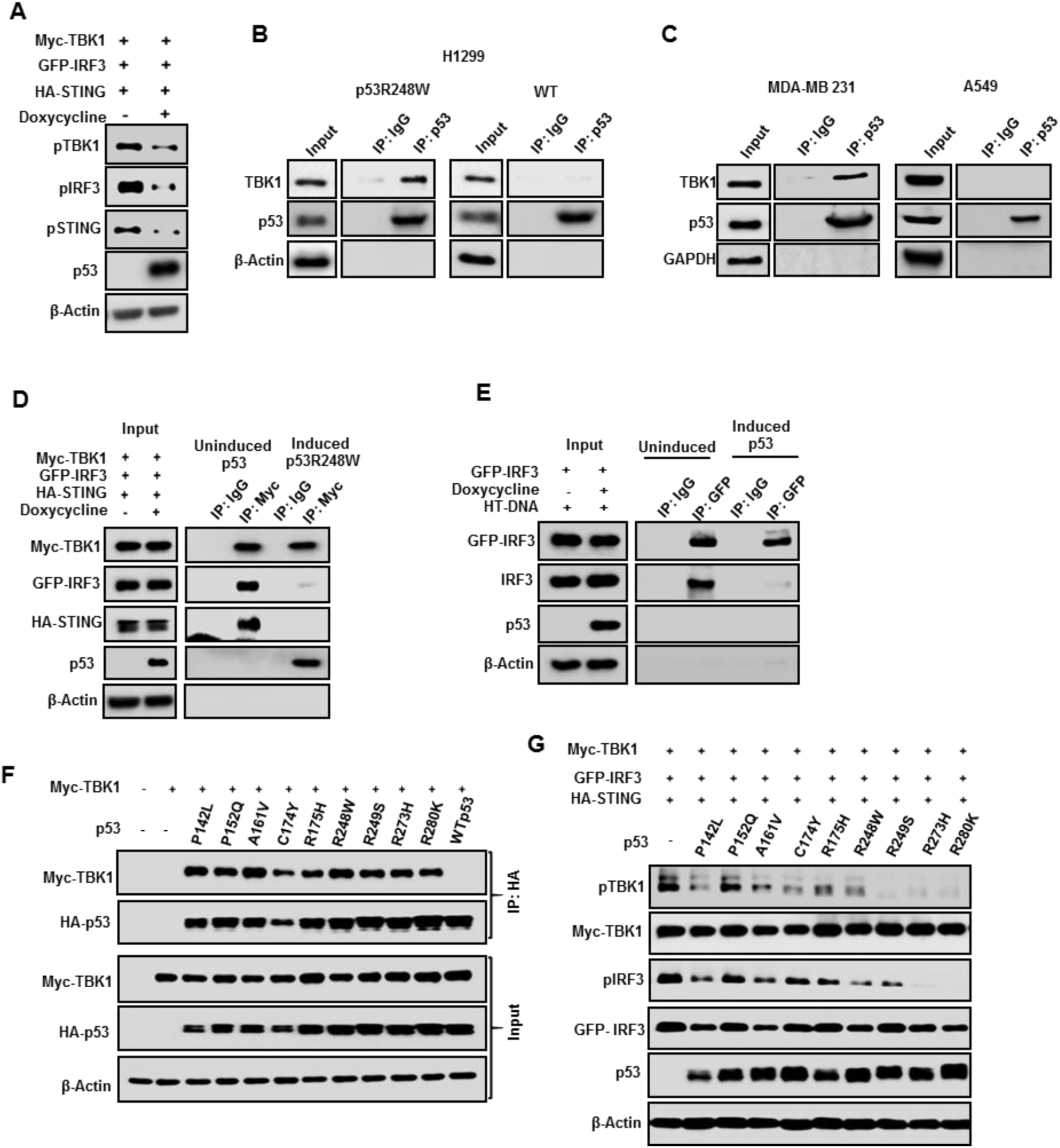
Mutant p53 interacts with TBK1 to prevent forming a trimeric complex with IRF3 and STING. (A) Myc-TBK1, GFP-IRF3 and HA-STING were co-transfected with or without p53R248W and subjected to western blot analysis. B) p53 null H1299 cells expressing inducible mutant p53R248W or WTp53 were induced with doxycycline treatment for 24 hrs and p53 was immunoprecipitated from the whole cell lysate. Lysates and immunoprecipitated (IP) were analyzed by SDS–PAGE and western blotting. (C) Endogenous mutant p53 (MDA-MB-231) or WTp53 (A549) was immunoprecipitated with p53 antibody. Cell Lysates and IP were analyzed by SDS–PAGE and western blot. (D) p53 null H1299 cells were left uninduced or induced with Doxycycline for 24 hrs to express p53R248W and co-transfected with Myc-TBK1, GFP-IRF3 and HA-STING. Cells were lysed and Myc-TBK1 was immunoprecipitated with Myc antibody. Cell Lysates and IP were analyzed by SDS–PAGE and western blot. (E) H1299 cells were induced to express p53R248W, transfected with GFP-IRF3 and subjected to HT-DNA treatment for 3 hrs. Cells were lysed and GFP-IRF3 was immunoprecipitated with GFP antibody. Whole cell lysate and IP were analyzed by SDS-PAGE and western blot. (F) In p53 null H1299 cells Myc-TBK1 was co-transfected with nine different mutant p53 and wtp53. Cells were lysed and p53 was immunoprecipitated. (G) H1299 cells were transfected with different mutant p53 with TBK1 and subjected to western blot analysis.

Our data with different cell lines carrying distinct p53 mutations suggested that both structural and DNA contact mtp53s could disable TBK1 (Figure 1). To compare abilities of different p53 mutants in an isogenic cellular context, we co-transfected TBK1 with different p53 mutants (P142L, P152Q, A161V, C174Y, R175H, R248W, R249S, R273H, R280K and WTp53) into H1299 cells. We observed that all the different mutants we tested were able to interact with TBK1, whereas WTp53 did not (Fig. 3F). Furthermore, the interaction of the mtp53 proteins with TBK1 also resulted in reduced phosphorylation of TBK1 and IRF3, albeit to varying degrees. In general, all the mutants reduced TBK1 substrate phosphorylation; notably, R249S, R273H and R280K almost completely suppressed TBK1 phosphorylation while R273H and R280K were the most potent suppressors of IRF3 phosphorylation (Fig. 3G). RT-PCR analysis of IFNB1 induction in cells co-transfected with TBK1/STING/IRF3 and the different mutants revealed that all the p53 mutants inhibited its induction by almost half (Fig. S5A). To map the sites of interaction between mtp53 and TBK1, we generated a series of p53 deletion mutants based on p53R249S and tested their ability to associate with TBK1. We found that deletion of amino acids 123-173 and 327-377 largely reduced the interaction between mtp53 and TBK1 (Fig. S5B). Taken together, our observations suggest that different p53 mutants can bind and prevent TBK1 from activating IRF3.

### Mtp53 tumors exhibit accelerated tumor growth in hosts with intact immune system

In addition to detecting and triggering a cell autonomous response to cytoplasmic DNA, the TBK1-STING-IRF3 pathway also serves to signal immune cells about the presence of tumor cells and thus functions in a non-cell autonomous manner.(*20, 30*) We therefore investigated whether mtp53 might modulate immune infiltration and impact tumor growth. Towards this end, we engineered 4T1 cells to express either PLVX or PLVX-p53R249S (referred as p53R249S hereafter) and used a syngeneic BALB/c mouse tumor model to assess tumor growth. Analysis of *in vitro* growth rates showed no difference between the PLVX and p53R249S cells (Fig. S6A). We injected 4T1 PLVX or p53R249S cells (5X10^4^) in the mammary fat pad of BALB/c mice. By 14 days post-inoculation, the p53R249S tumors grew much faster than the PLVX ones and by the end of the experiment, they were approximately twice as large (Fig. 4A, 4B, and 4C). We observed no significant difference in immunohistochemical detection of the proliferation marker, Ki67, between PLVX and p53R249S tumors (Fig. S6B). These data indicated that the increased growth of the p53R249S tumors was not due to an intrinsic proliferative advantage and suggested that cell-extrinsic signaling underlie these differences. To directly assess if the immune system played a role in the different tumor growth rates, we inoculated the same 4T1 PLVX and p53R249S cells into NOD/SCID mice and monitored tumor growth. In this immunocompromised mouse, we failed to find any differences in the growth rates of the tumors over the course of the experiment (Fig. 4D, 4E and Fig. S6C). We also assessed tumor vascularization by immunohistochemical detection of the endothelial marker, CD31. Surprisingly, in the tumors grown in the immunocompetent BALB/c mouse model, we found higher levels of CD31 positive cells in the p53R249S tumors than the PLVX. In contrast, tumors grown in NOD/SCID mice did not have any apparent difference in CD31 staining (Fig. 4F, and 4G). Thus, p53R249S exhibits gain-of-function activities of increased tumor growth and neoangiogenesis *in vivo* only in hosts with an intact immune system.

**Figure 4:**
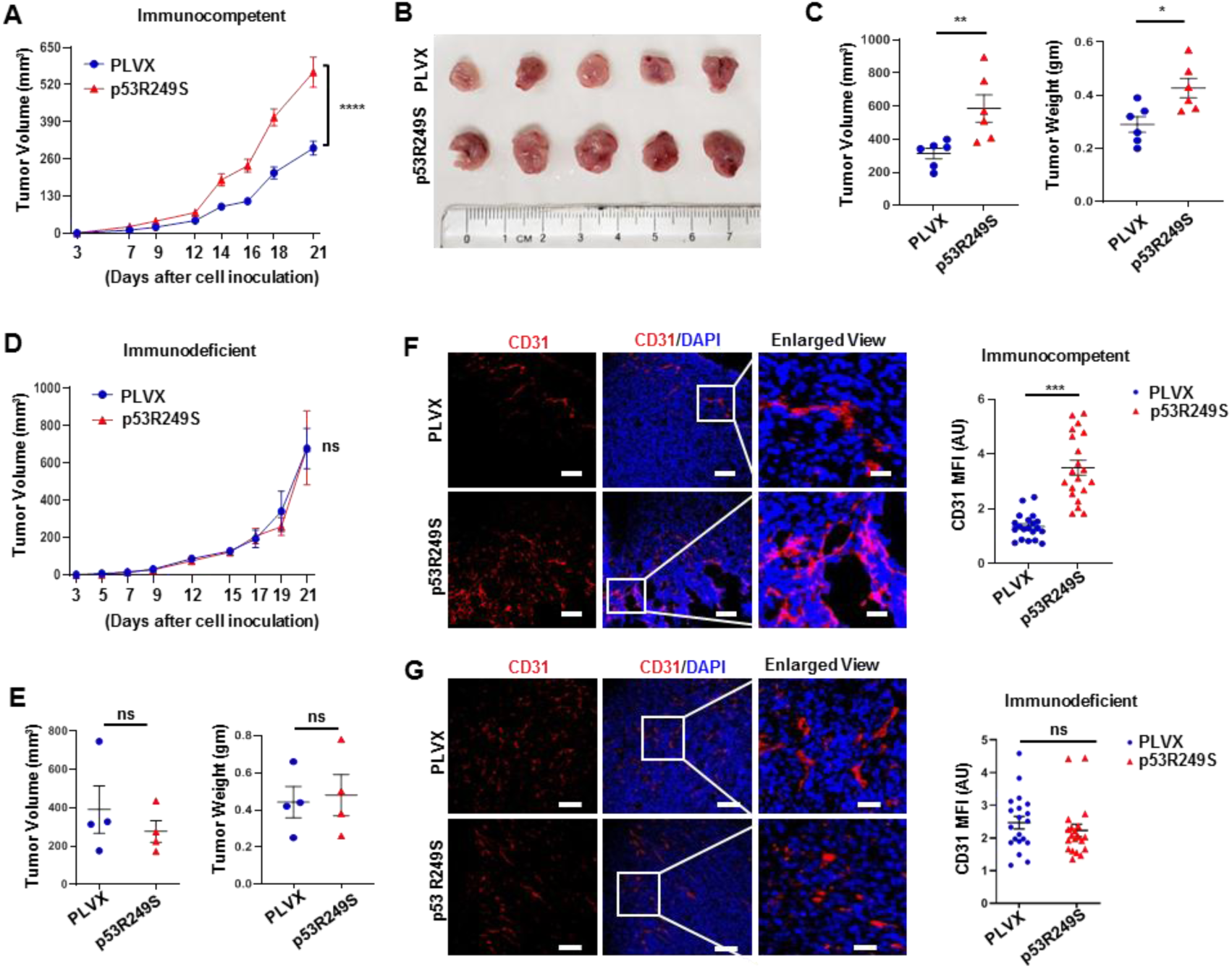
Mtp53 tumors exhibit accelerated tumor growth in hosts with intact immune system. (A) 5 x 104 4T1 cells expressing PLVX or p53R249S was injected onto the mammary gland of immunocompetent female BALB/c mice (n=10). All mice were sacked on day 21 and graphical quantification represents the tumor growth rate in mice. (B) Representative image showed tumor volume difference in BALB/c mice and (C) Graphical quantification of difference in tumor volume and weight on day 21 in PLVX and p53R249S cohorts (n= 6). (D) 5 x 104 4T1 cells expressing PLVX or p53R249S was injected onto the mammary gland of immunodeficient NOD/SCID mice (n=4). All mice were sacked on day 21 and graphical quantification represents the tumor growth rate in NOD/SCID mice. (E) Graphical quantification of difference in tumor volume and weight in PLVX and p53R249S cohorts in NOD/SCID mice. Representative confocal micrographs of 4T1 tumor sections from (F) immunocompetent (BALB/c) and (G) immunodeficient (NOD/SCID) mice stained with angiogenic marker CD31. (Right) Representative quantification graphs of endothelial marker CD31 intensity expression in PLVX and p53R249S tumors grown in either immunocompetent or immunodeficient mice (n=20) respectively. Quantification graphs: FoV= Field of View, In all panels, error bars represent mean with standard error. In scatter dot plots, each dots represent one mice, p values are based on Student’s t test. ∗∗∗p < 0.001, ∗∗p < 0.01, ∗p < 0.05, ns=non-significant.

### Mutant p53 suppresses immune surveillance to support tumor growth *in vivo*

To gain mechanistic insight into the phenotypic differences between these tumors, we did western blot analysis of 4T1 PLVX or p53R249S tumor lysates to detect IRF3 phosphorylation and RT-PCR analysis to assess IFNB1 mRNA levels. We observed reduced phosphorylation of IRF3 in the p53R249S tumors (Fig. 5A). Importantly, comparison of IFNB1 mRNA levels demonstrated that p53R249S tumors had reduced production of this cytokine (Fig. 5B). Taken together, our results suggest that mutants p53 suppressed IRF3 activation and IFNB1 production *in vivo*.

**Figure 5:**
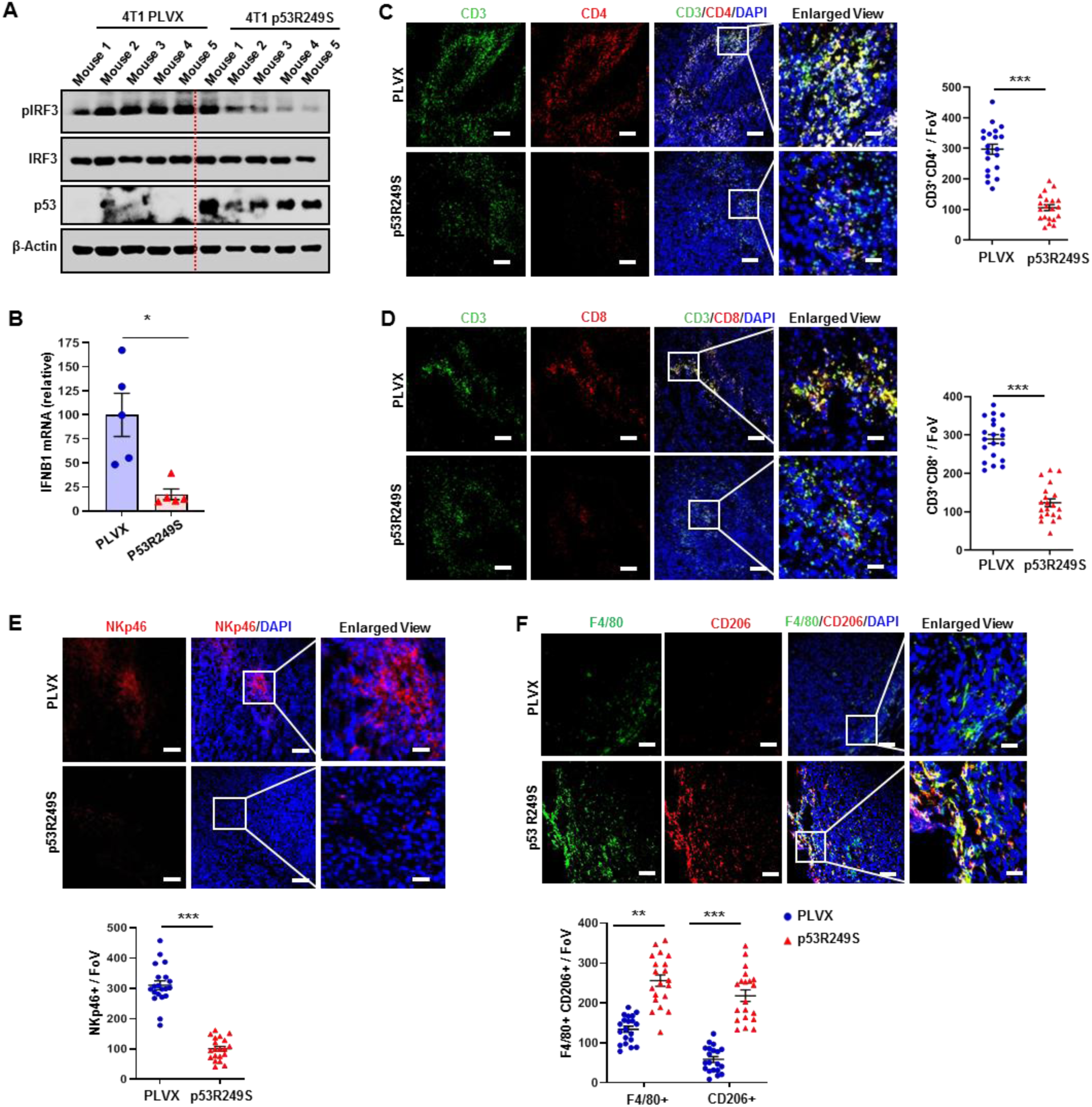
Mutant p53 suppresses immune surveillance to support tumor growth *in vivo*. 4T1 tumors in BALB/c mice were resected on day 21, cut into pieces (A) Lysed with RIPA and subjected to western blot analysis. (B) RNA was isolated and subjected to RT-PCR for IFNB1. Representative confocal micrographs of (C) CD3+CD4+ T-helper and (D) CD3+CD8+ T-cytotoxic lymphocyte infiltration. (Right) Representative graphs showed quantification of CD3+CD4+ T-helper and CD3+CD8+ T-cytotoxic lymphocyte (n=20) respectively. (E) Representative confocal micrographs of expression of NKp46, NK T-cells in 4T1 PLVX and p53R249S tumor sections. (Bottom) representative graph shows quantification of NK T-cells recruitment (n=20) in 4T1 PLVX and p53R249S tumor sections. (F) Representative confocal images depicting the F4/80+CD206+ M2 type of TAMs in PLVX and p53R249S expressing tumors isolated from BALB/c mice on Day 21. (Bottom) Representative graph indicates quantitation of F4/80+/CD206+ TAMs in PLVX and p53R249S tumor sections (n=20). Images were captured at 20X, scale bar 25 μm except in enlarged panel which is 100 μm. Quantification graphs: In all panels, error bars represent mean with standard error. p values are based on Student’s t test. ***p < 0.001, ∗p < 0.05, ns=non-significant.

Previous reports have shown that the innate immune STING-TBK1-IRF3 pathway plays an important role in anti-cancer immunity *in vivo* via activation of the type I interferon (IFN) response.(*20, 30, 32*) Functionally, cancer cell production of type I IFN enhances recruitment of natural killer (NK) cells, T lymphocytes, and macrophages.(*20, 30, 32*) Cytotoxic T lymphocytes, NK cells and macrophages play major roles in controlling tumor growth and thus we decided to assess if mtp53 modulated the immune infiltration within the tumor microenvironment. Whereas PLVX tumors had an abundance of both CD4+ and CD8+ T-lymphocytes, the p53R249S tumors showed relatively less infiltration by these cells. (Fig. 5C and 5D). Furthermore, we observed a robust reduction of NK cell infiltration in the tumors expressing p53R249S compared to its PLVX counterpart (Fig. 5E). Consistent with the reduced cytotoxic T-lymphocytes and NK cells infiltration in p53R249S tumors, we detected reduced apoptotic cells in these tumors relative to its PLVX counterpart (Fig. S7A). Conversely, macrophages detected by F4/80^+^ staining were more abundant in the p53R249S tumors as determined by immunohistochemistry. Since TAM polarization to the M2 state has been shown to have a pro-tumorigenic role by promoting neoangiogenesis, we assessed the F4/80+ macrophages for expression of CD206, a widely used marker for the macrophage M2 polarized state. This analysis indicated a stark difference between the PLVX and p53R249S tumors. Whereas most of the TAMs in PLVX tumors were negative for CD206 staining, the majority of them were in the M2 polarized state (F4/80+CD206+ dual positive) in the p53R249S tumors (Fig. 5F). Given the lack of enhanced growth of the p53R249S tumors relative to the PLVX ones in the NOD/SCID mice, we performed IHC to assess the degree of macrophage infiltration and polarization (Fig. S7B). Consistent with the known defective function of macrophages in NOD/SCID mice, we observed a 10-fold decrease in macrophage infiltration in both PLVX and p53R249S tumors, as well as a virtually complete absence of CD206 positive M2 cells.(*33, 34*) Our results reveal that mtp53 promotes a pro-tumor microenvironment by reducing tumor infiltration of critical mediators of the anti-tumor response (CD8+ and NK cells) and promoting the phenotypic alteration of immune cells (M2 macrophages) to provide support for tumor growth.

We were intrigued to find such a striking difference in the presence M2 polarized macrophages between the tumors. Since the STING/TBK1/IRF3 pathway is critical for IFNB1 expression, we considered the possibility that reduced IFNB1 production in p53R249S tumors resulted in increased macrophage polarization to the M2 state. The latter is consistent with a previously reported function of IFNB1, which is to maintain cells in the M1 phenotype.(*36, 37*) Therefore, to test if differences in cytokine production between the PLVX and p53R249S cells impacted macrophage polarization, we cultured RAW264.7 macrophage cells with the 4T1 PLVX or p53R249S conditioned medium. Macrophages cultured for 24 hours with PLVX conditioned medium expressed more mRNA of the M1 macrophage markers, TNFα and CD86, than did macrophages cultured in p53R249S conditioned medium. In contrast, macrophages exposed to the p53R249S conditioned medium exhibited higher expression of the M2 marker mRNA, IL-10 (Fig. S8A). To determine if IFNB1 was reinforcing the M1 phenotype in the macrophages, we repeated the experiment with PLVX conditioned medium alone or the same medium in which we had added IFNB1 neutralizing antibody. Incubation of the macrophages with medium containing the IFNB1 neutralizing antibody no longer retained the M1 phenotype as determined by their reduced expression of the M1 marker, TNFα (Fig. S8B). This observation reinforced the notion that IFNB1 plays a critical role in the maintenance of the M1 phenotype. Next we considered that supplementing the p53R249S conditioned medium with IFNB1 protein might prevent M2 macrophage polarization. We had previously determined that the PLVX cells secreted approximately 15 pg/ml of IFNB1, and thus we supplemented the p53R249S medium with additional IFNB1 to achieve this concentration. Addition of IFNB1 to the p53R249S conditioned medium resulted in reduced expression of the M2 marker, IL-10 (Fig. S8C). Collectively our data indicate that by blocking type I interferon (IFNB1) production, mtp53 drives tumorigenesis by modulating immune cell recruitment and macrophage polarization to support tumor growth.

### Mutant p53 suppresses immune surveillance to support KPC tumor growth

To further substantiate our assertion that mtp53 altered the tumor microenvironment to promote favorable conditions for tumor growth, we used KPC cells with inducible p53 shRNA to generate tumors in a syngeneic tumor host. KPC cells infected with a doxycycline inducible empty vector (EV) or p53 shRNA (shp53) were treated *in vitro* with doxycycline for 2 days, and then 1X10^5^ cells were injection into the dorsal lateral side of C57BL/6 mice. All the mice were given doxycycline (20 mg/kg) by oral gavage. Strikingly, mtp53 knockdown strongly reduced the tumor growth rate (Fig. 6A, 6B and 6C). Similar to the 4T1-mtp53 expressing cells, KPC tumors expressing mtp53 (EV) were highly angiogenic as assessed by IHC staining for CD31 (Fig. 6D). Tumors from the shp53 cells exhibited strongly reduced staining for CD31, indicating that loss of mtp53 compromised their angiogenic potential. Previously it was reported that mtp53 knockdown in KPC cells had no effect on primary tumor growth in immune-deficient mice, which led us to hypothesize that the reduced tumor growth observed in the immune-proficient syngeneic host was due to altered immune cell recruitment.(*35*) Analysis of T-lymphocytes in the tumors revealed that mtp53 knockdown resulted in robust infiltration of CD3+, CD4+, CD8+ cells and NK cells (Fig. 6E, 6F, 6G and S9A, S9B, S9C). Remarkably, mtp53 knockdown also resulted in a reduction of M2-polarized macrophages in the tumors (Fig. 6H and S9D). Taken together, our data support the notion that mtp53 promotes tumorigenesis by generating a tumor-supportive microenvironment.

**Figure 6:**
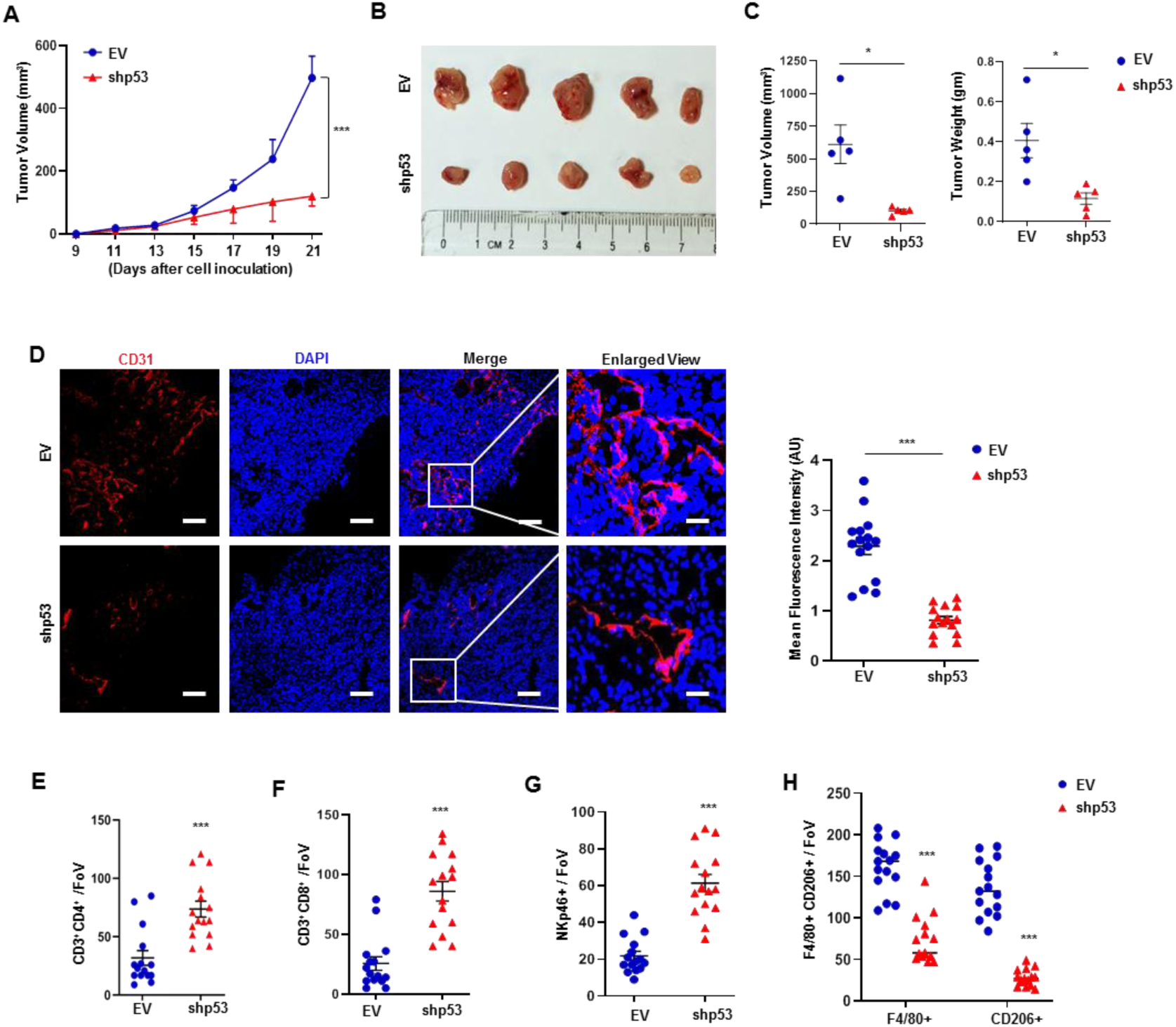
Mutant p53 suppresses immune surveillance to support KPC tumor growth. (A) KPC inducible EV/ shp53 (1×105) cells were injected (subcutaneous) in the male C57BL/6 mice. Doxycycline (20 mg/kg) was given orally every other day to all the mice to induced either EV or shp53 starting from day 4. Tumor volume was monitored and measured manually using slide calipers. (B) All KPC tumor harboring mice were sacked on day 21 and representative image showed tumor volume difference between EV and shp53. (C) Representative graphical quantification of difference in tumor volume and weight on day 21 in EV and shp53 KPC tumor cohorts (n= 5). (D) Representative confocal micrographs of EV and shp53 KPC tumor sections stained with angiogenic marker CD31. Graph at the right indicates quantitation of mean fluorescence intensity of CD31 in EV and shp53 tumors. Representative graphs showed quantification of (E) CD3+CD4+ T-helper and (F) CD3+CD8+ T-cytotoxic lymphocyte (G) NK T-cells and (H) F4/80+/CD206+ TAMs in EV and shp53 tumor sections (n=15). Images were captured at 20X, scale bar 25 μm except in enlarged panel which is 100 μm. Quantification graphs: In all panels, error bars represent mean with standard error. In scatter dot plots, each dots represent one mice, p values are based on Student’s t test. ***p < 0.001, ∗p < 0.05, ns=non-significant.

### Ectopic TBK1 expression overrides mtp53’s pro-tumorigenic immunomodulatory activity

Our overall findings indicated that mtp53 suppresses TBK1 function and thereby prevents downstream signaling to IRF3, resulting in deficient paracrine signaling to immune cells. Therefore, we reasoned that ectopic TBK1 expression might restore signaling to IRF3. To test this possibility, we infected BT549 and MDA-MB-231 cells with an empty or TBK1 expressing vector. TBK1 overexpression was sufficient to restore IRF3 (and STING) phosphorylation and induced the expression of multiple IRF3 target genes in both cell lines (Fig. S10A and S10B). Next, to test the consequence of restoring IRF3 activation on tumor growth, we engineered the 4T1-PLVX and 4T1-p53R249S cells to express a doxycycline inducible TBK1 and characterized these cells *in vitro* (Fig. S11A). TBK1 induction resulted in an approximately 2-fold increase in TBK1 protein levels, but this did not affect cell growth as assayed by MTT analysis (Fig. S11B). TBK1 induction correlated with a 4-fold increase in IFNB1 mRNA in PLVX cells and a 2-fold increase in p53R249S cells (Fig. S11C). Importantly, ELISA-based measurement of IFNB1 released in conditioned medium showed that TBK1 induction in the p53R249S cells restored IFNB1 to approximately equivalent levels to those detected in the PLVX cells (Fig. S11D). Next, we sought to determine whether induction of TBK1 would alter the conditioned medium of p53R249S resulting in reduced RAW264.7 macrophage polarization *in vitro*. Exposure of the RAW264.7 cells to the conditioned medium from the PLVX cells with TBK1 induction further reinforced the M1 phenotype as indicated by increased expression of M1 markers by the macrophages. The conditioned medium from the p53R249S cells with induced TBK1 had a moderately diminished capacity to promote the M2 phenotype in the macrophages since their TNFα and IL-10 mRNA expression was reduced (Fig. S11E).

We next tested whether TBK1 induction would be sufficient to impact tumor growth in the BALB/c syngeneic mice model. Female BALB/c mice were injected with 4T1 PLVX or p53R249S (5X10^4^) doxycycline inducible TBK1 cells. Mice were given doxycycline orally according to the schedule and tumor size was monitored (Fig. 7A). Indeed, TBK1 induction strongly reduced the growth rate of the p53R249S tumors to the extent that at the end of the experiment these tumors were smaller than the PLVX tumors. We also observed that TBK1 expression in PLVX tumors modestly affected tumor growth (Fig. 7B and Fig.S11F). We did RT-PCR for mRNA expression in the TBK1 induced tumors. Our results suggests a 4-fold IFNB1 mRNA induction in p53R249S induced TBK1 tumor (Fig. 7C). Strikingly, TBK1 induction in the p53R249S tumors reduced CD31 immunohistochemical staining, indicating that neovascularization was blocked (Fig.S11G). Furthermore, assessment of lymphocyte infiltration in response to TBK1 overexpression in p53R249S tumors revealed a robust recruitment of CD4^+^ T-helper and CD8^+^ T-cytotoxic lymphocytes as well as an increase in NK cell infiltration of these tumors (Fig. 7D, 7E, 7F, 7G and Fig.S12 and S13A). Moreover, TBK1 overexpression in p53R249S tumors resulted in a reduction in M2 polarized macrophages (Fig. 7H and Fig. S13B). Taken together, our data suggests that the ability of mtp53 to foster a pro-tumorigenic micro-environment can be reversed by restoring TBK1 signaling.

**Figure 7:**
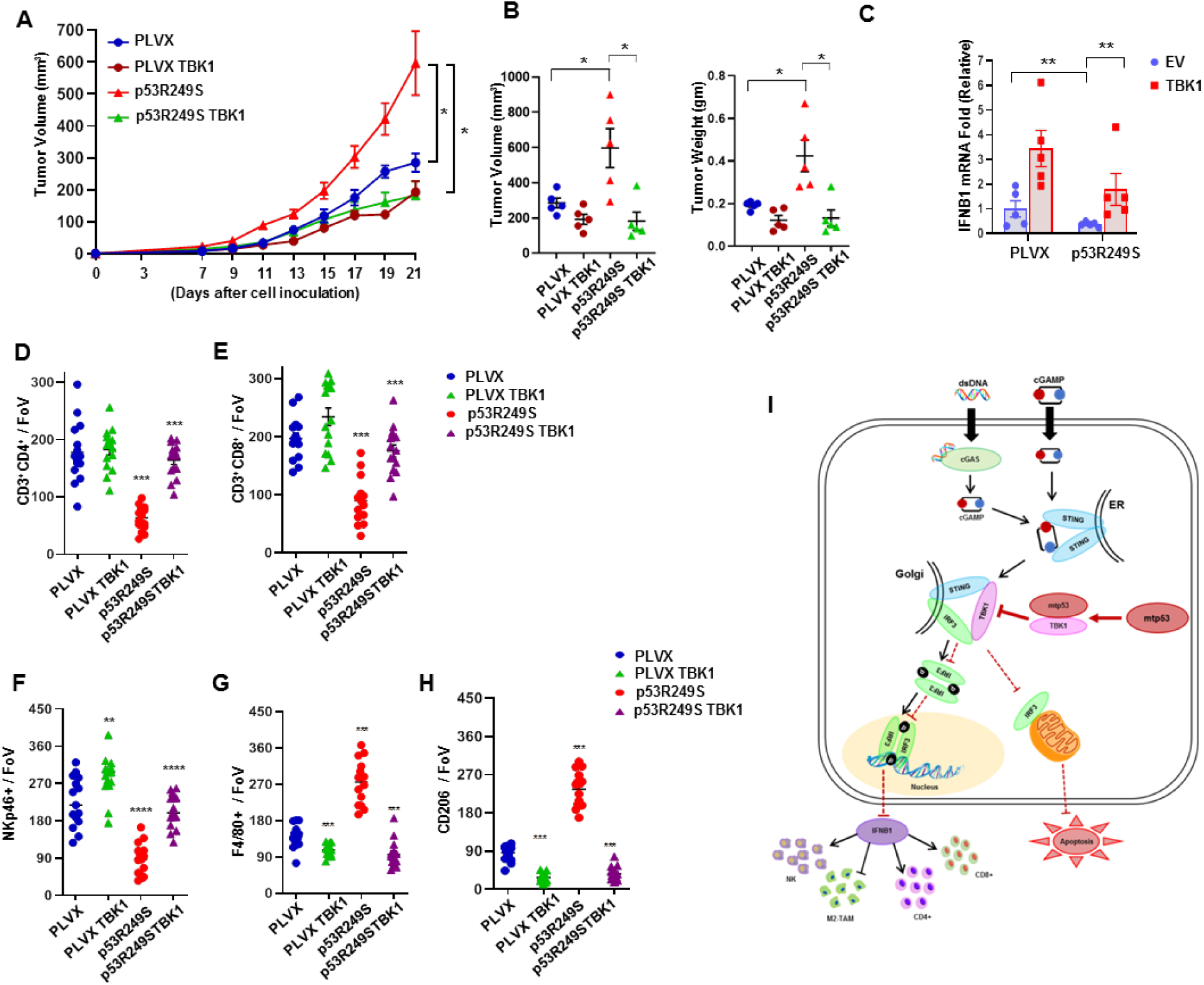
Ectopic TBK1 expression overrides mutant p53’s effect. (A) 5 x 10^4^ PLVX or p53R249S inducible EV or TBK1 4T1 cells were injected in the mammary fat pad of female BALB/c mice. Doxycycline was administered from Day 5 and tumor volume was measured. (B) Graphical quantification showing the tumor volumes and weight differences of different cohorts. (C) Mice were sacrificed on Day 21; tumors were excised, and RNA was isolated. Representative graph indicate quantitative mRNA expression of IFNB1 in EV or TBK1 inducible 4T1 PLVX or p53R249S tumors. (D-H) Representative quantitation from the cryo-section of different tumor cohort showed CD3^+^CD4^+^ T-helper cells, CD3^+^CD8^+^ T-cytotoxic subsets, NKp46^+^ NK cells and F480+, CD206+ M2 macrophage subsets in PLVX or p53R249S tumors induced TBK1 in BALB/c mice (n=15). (I) Graphical representation of mutant p53 disabling the innate immune response signaling pathway. Mutant p53 expressing cells fail to activate the type I interferon response to modulate tumor microenvironment (non cell autonomous) and also suppress mitochondria mediated apoptosis (cell intrinsic). Quantification graphs: In all panels, error bars represent mean with standard error. In scatter dot plots, each dots represent one mice, p values are based on Student’s t test. ^∗∗∗^p < 0.001, ^∗∗^p < 0.01, ^∗^p < 0.05, ns=non-significant.

## Discussion

Mutation of the TP53 gene is one of the most frequent genetic lesions in human cancers. Although mutant p53 proteins exhibit gain-of-function activities that contribute to tumor development and progression, recent studies have indicated that such oncogenic functions of mtp53 are context dependent.(*38, 39*) Importantly, hotspot mtp53 mutants were found to lack the ability to confer a growth advantage *in vitro*.(*38, 39*) However, since our study is the initial report of this novel gain-of-function activity of mtp53, this was not assessed by those studies. An additional caveat to consider is that some cell lines have lost cGAS/STING signaling and thus it would not be possible to assess this mtp53 gain-of-function in those cells.(*40–43*)

In our study, we utilized multiple cell lines (human and mouse), multiple approaches (shRNA, ectopic expression, MEFs from genetically engineered mice) to corroborate the novel gain-of-function activity of mtp53. Therefore, our data provide a novel mechanism by which mtp53 can interact with TBK1 and suppress downstream IRF3 activation to exert GOF that compromises cell intrinsic (apoptosis) and extrinsic (immunomodulatory) anti-tumor activities of the STING-TBK1-IRF3 pathway. We observed that different p53 mutants were able to suppress this pathway. This raises the possibility that p53 mutants that potently interfere with the pathway will be selected for and overrepresented in cancer. Thus, suppression of the innate immune pathway may be a selective mechanism that drives the occurrence of hotspot p53 mutations.

In direct contrast to mtp53, our data suggest that wildtype p53 contributes to the activation of the cGAS/STING pathway. The precise mechanism underlying this remains to be determined but it is likely that IFI16, a transcriptional target of wildtype p53, is involved. (*26–28*) Interestingly, wildtype p53 is transcriptionally induced by and enhances interferon signaling.(*44, 45*) Nevertheless, since wildtype p53 does not interact with TBK1, this indicates that mtp53 controls the innate immune pathway through a distinct mechanism.

Intriguingly, mtp53 proteins are tumor specific neo-antigens that are immunogenic and would be predicted to render cancer cells susceptible to immune editing. (*46–49*) However, despite eliciting immunogenic responses in tumor infiltrating lymphocytes, mtp53 expressing cells persist. We observed that mtp53 altered immune cell infiltration/phenotypes and this could be reversed by reactivating IRF3 through TBK1 overexpression. We speculate that suppression of the STING-TBK1-IRF3 pathway by mtp53 renders tumors immunologically “cold”, thereby permitting cancer cells that express this neo-antigen to evade immune detection. Importantly, disruption of the mtp53/TBK1 complex switches the tumor microenvironment from cold to hot and permits the immune system to limit tumor growth (Fig. 7I). Therapeutic approaches aimed at activating TBK1 function could restore tumor suppressing immune-surveillance and eliminate mutant p53 expressing tumors.

## Acknowledgements

This work was supported by NCI CA166974-01A1, the NY State Empire Investment Program, the Stony Brook Renaissance School of Medicine and Cancer Center, the TRO Carol M. Baldwin Award, the Stony Brook Cancer Center Bahl IDEA Award and the Lynn November Pilot Funds for Therapeutic Development in Aggressive Breast Cancer from the Stony Brook Cancer Center. We thank Dr. Richard Linn (Stony Brook University, NY, USA) for sharing KPC cells and Dr. Nancy C. Reich (Stony Brook University, NY, USA) for providing us with the pcDNA3GFP-IRF3 clone. We would like to acknowledge the technical support provided by the Research Flow cytometry core facility, Department of Pathology, Stony Brook Renaissance School of Medicine.

## Authors Contributions

L.A.M. conceived and supervised the project. L.A.M. and M.G. designed the research. M.G. performed most of the experiments, L.A.M. assisted in experiments. S.S. helped in all the animal experiments and statistical analysis. J.B. helped in animal experiments. R. N. helped in cloning the plasmids and assisted in experiments. A.P. and T.I. isolated the MEFs and edited manuscript. L.A.M and M.G. analyzed the data and wrote the manuscript. A.W.M.V. helped in animal experiments and edited the manuscript.

---

**Figure S1:**
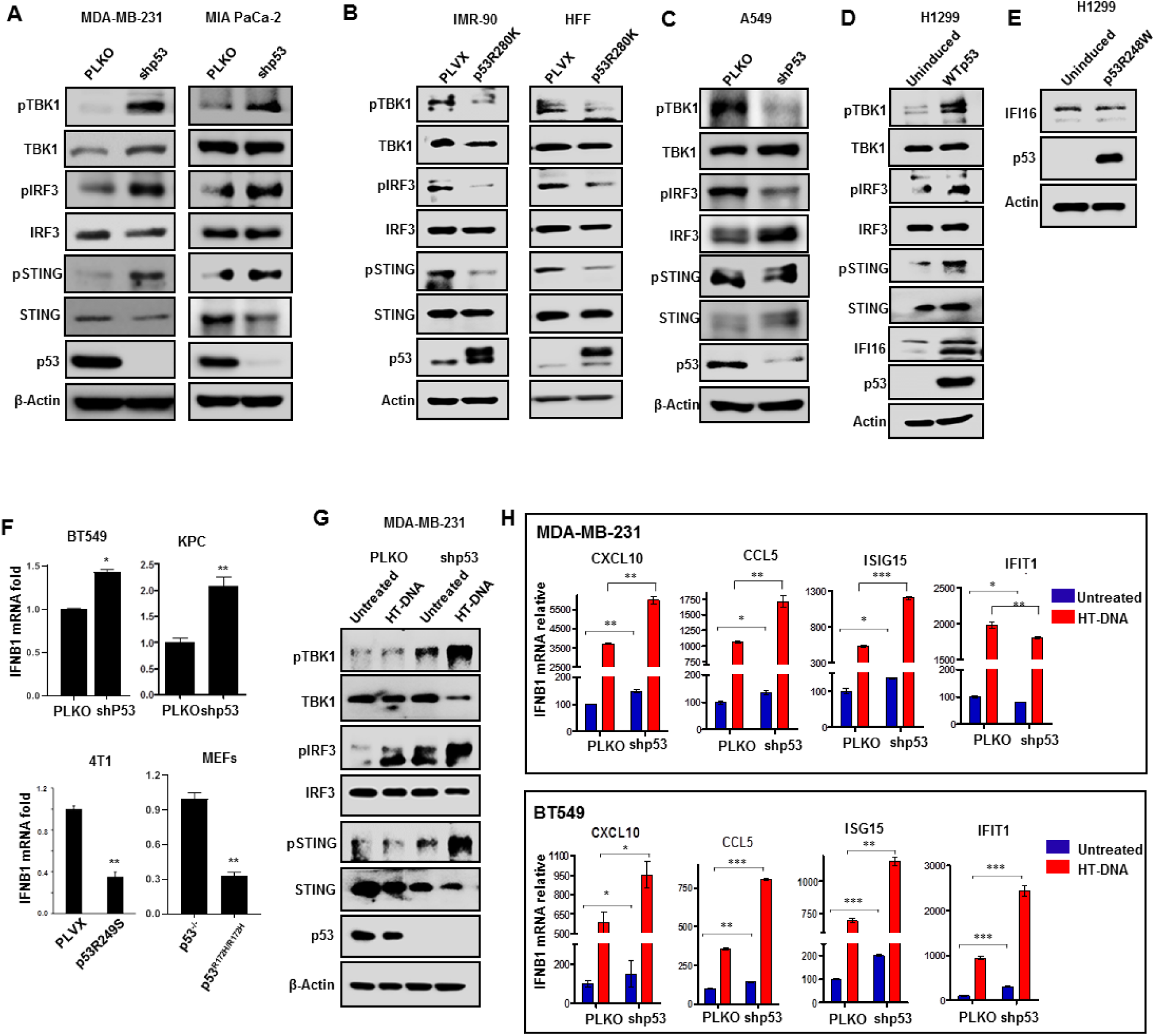
Mutant p53 suppresses activation of innate immune pathway. (A) Indicated cells were infected with virus containing control or p53 targeting shRNA and subjected to Immunoblot analysis. (B) “Non-cancerous” human fibroblasts IMR-90 and HFF cells were engineered to express p53R280K and subjected to western blotting. (C) A549 cells were infected with virus containing EV (PLKO) or p53shRNA and subjected to western blot. P53 null H1299 cells were induced with doxycycline express either (D) WTp53 or (E) mutant p53R248W and subjected to western blot. (F) Representative RT-PCR data showing mutant p53 attenuates IFNB1 mRNA in the designated cells. (G) Indicated PLKO or shp53 cells were treated with 2 μg/ml of HT-DNA for 3 hrs, cells were harvested and subjected to western blot analysis. (H) p53 knock down BT549 or MDA-MB-231 cells were treated with HT-DNA for 18 hrs, cells were harvested, RNA was isolated from the cells for RT-PCR. Quantification graphs: In all panels, error bars represent mean with standard deviation. p values are based on Student’s t test. ^∗∗∗^p < 0.001, ^∗∗^p < 0.01, ^∗^p < 0.05, ns=non-significant.

**Figure S2:**
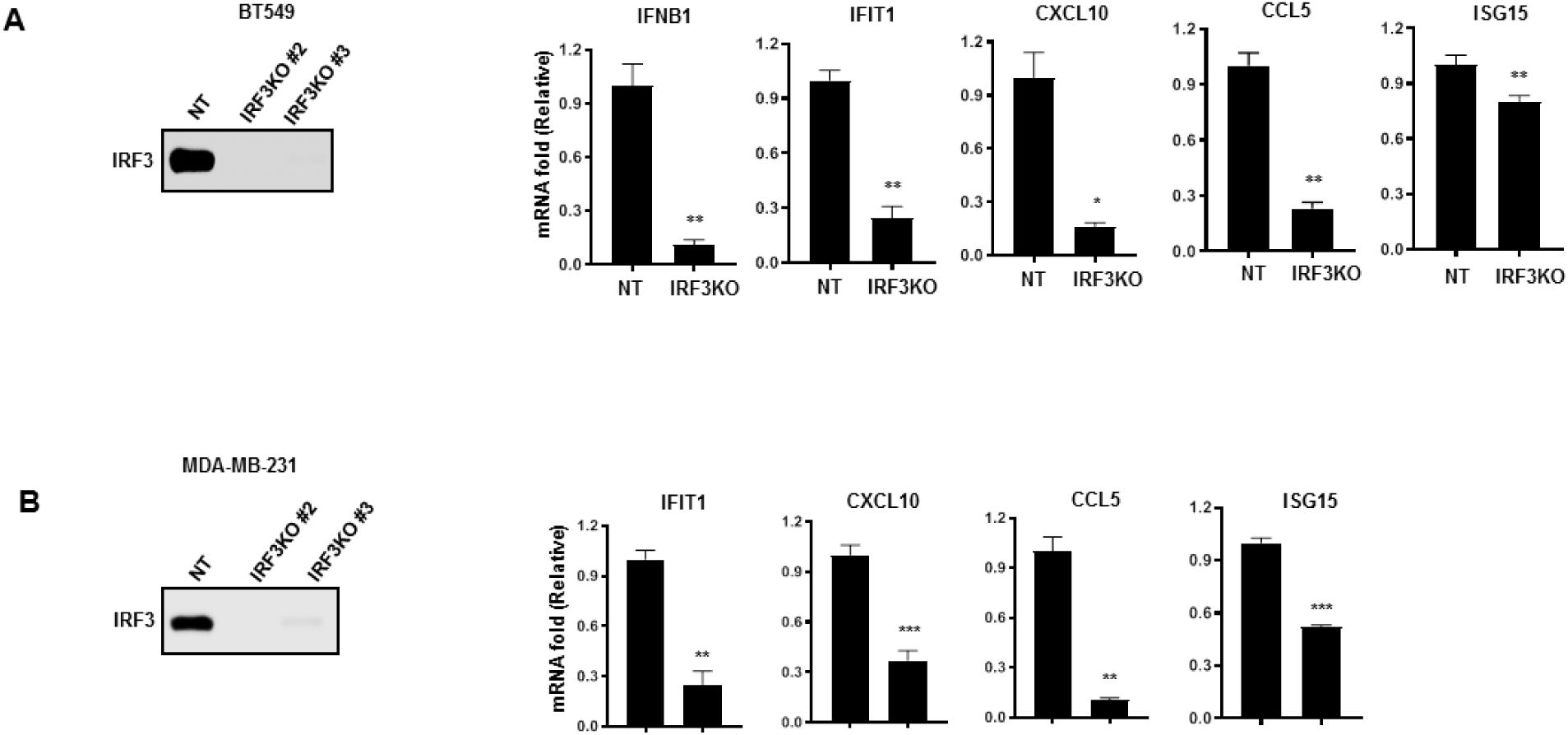
IRF3 target chemokines. IRF3 knock out cells were prepared using CRISPR-Cas9 technique and cells were subjected to western blot and RT-PCR analysis. Graph showed mRNA expression of CXCL10, CCL5, ISG15, IFIT1 in IRF3KO #2 (A) BT549 and (B) MDA-MB-231 cells. Quantification graphs: In all panels, error bars represent mean with standard deviation. p values are based on Student’s t test. ^∗∗∗^p < 0.001, ^∗∗^p < 0.01, ^∗^p < 0.05.

**Figure S3:**
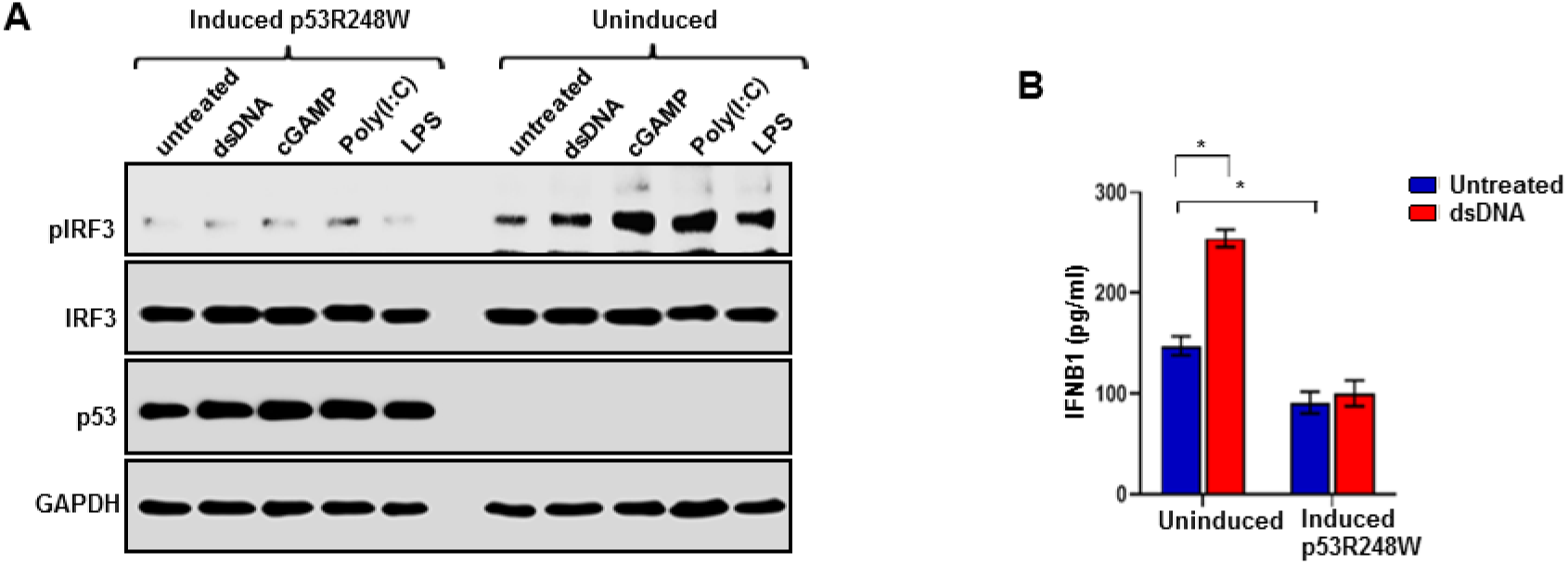
Mutant p53 blocks IRF3 activation. (A) H1299 cells were induced to express p53R248W and treated with dsDNA, cGAMP, poly(I:C) or LPS for 3 hrs and subjected to western blot. (B) Representative graph shows ELISA data of IFNB1 released from H1299 cells uninduced and induced p53R248W and treated with 2 μg/ml dsDNA for 18 hrs. Quantification graphs: In all panels, error bars represent mean with standard deviation. p values are based on Student’s t test. ^∗∗∗^p < 0.001, ^∗∗^p < 0.01, ^∗^p < 0.05.

**Figure S4:**
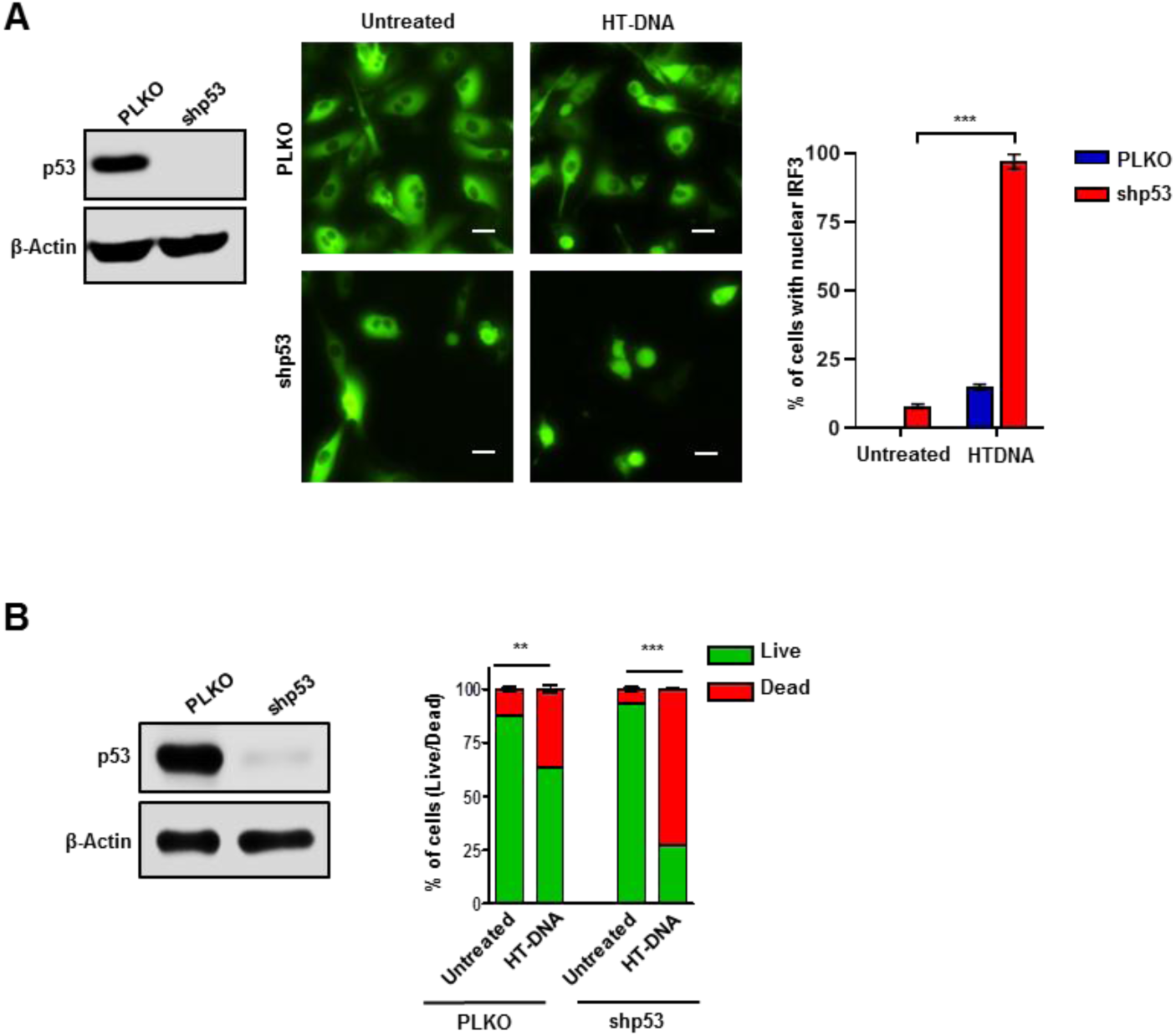
Mutant p53 blocks IRF3 nuclear translocation and attenuates IRF3 mediated apoptosis. (A) Representative fluorescent images depicting nuclear translocation of GFP-IRF3 in PLKO or mutant p53 knock down MDA-MB-231 cells. Cells were treated with 2µg/ml HT-DNA for 3 hrs and IRF3 localization was visualized. Representative graph shows quantification of percent of cells with nuclear IRF3. Images were captured in Nikon T*i* fluorescence microscopy at 60X magnification. Scale bar 20 µm. (B) Graphical representation of apoptosis quantification in PLKO or p53KD MDA-MB-231 cells that were treated with 2ug/ml of HT-DNA for 24 hrs using flow cytometry. Immunoblots showing p53 knock down efficiency. Scale bar 20μm. Images were captured in a Nikon T*i* inverted fluorescence microscope at 60X magnification. Quantification graphs: FoV= Field of View, (n=20) In all panels, error bars represent mean with standard deviation. p values are based on Student’s t test. ^∗∗∗^p < 0.001, ^∗∗^p < 0.01.

**Figure S5:**
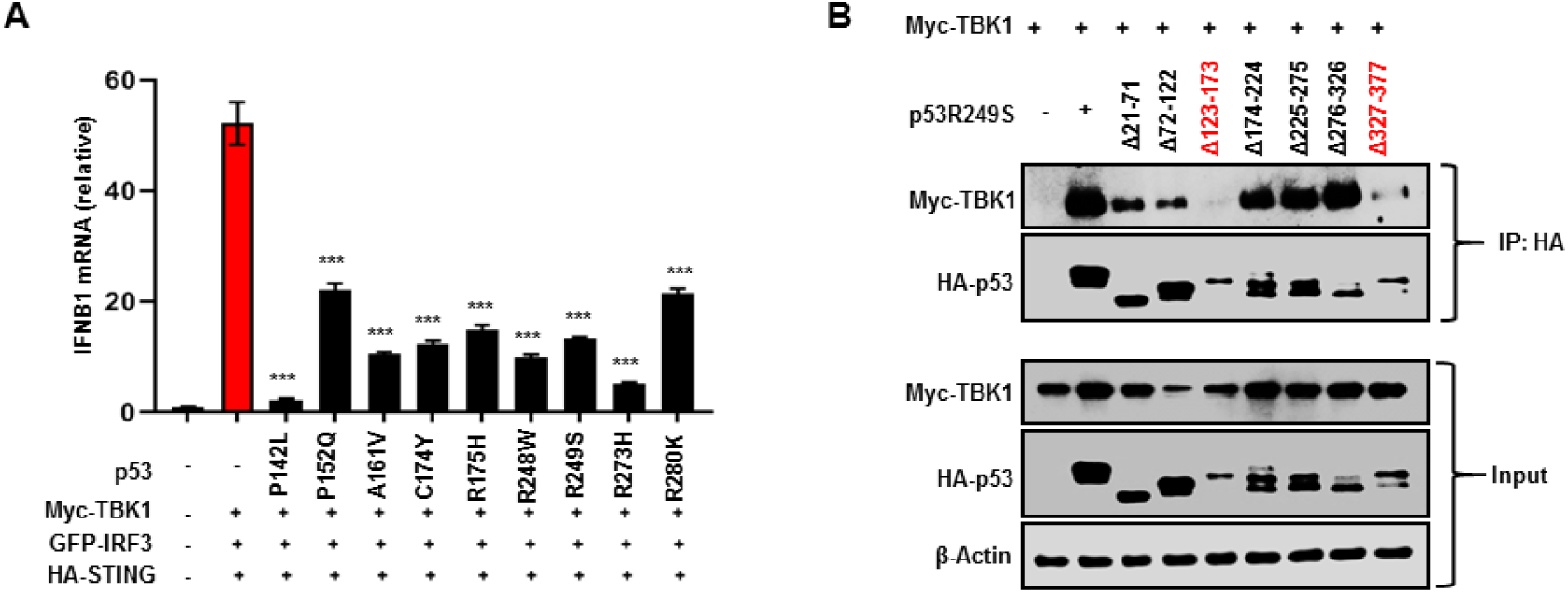
p53 mutants suppresses IRF3 mediated IFNB1 induction. (A) Quantification of IFNB1 mRNA expression in p53 null H1299 cells that were transfected with Myc-TBK1, GFP-IRF3 and HA-STING in absence or presence different p53 mutants. Cells were harvested after 24 hrs, and RT-PCR for IFNB1 was performed. (B) H1299 cells were transfected with Myc-TBK1 and seven different deletion mutants of HA-p53R249S. Cells were lysed and mutant p53 was immunoprecipitated using HA antibody. Quantification graphs: In all panels, error bars represent mean with standard deviation. p values are based on Student’s t test. ^∗∗∗^p < 0.001, ^∗∗^p < 0.01.

**Figure S6:**
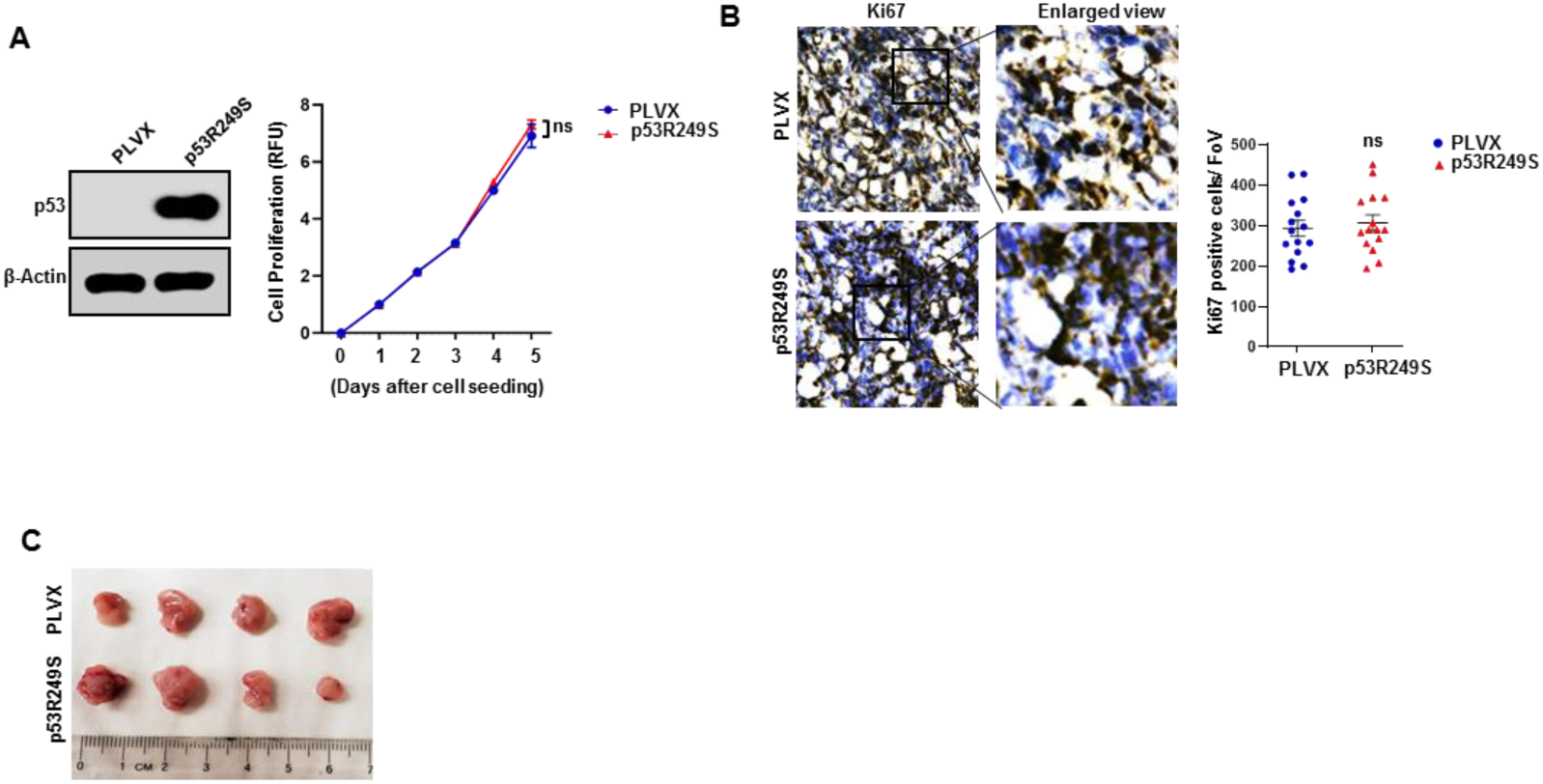
p53 mutation has no significant effect in tumor cell proliferation *in vitro* but *in vivo* mutant p53 strongly supports tumor growth. (A) Immunoblots shows p53 null 4T1 cells were engineered to express PLVX or p53R249S. Representative graphical quantification of *in vitro* cell proliferation between 4T1 PLVX and p53R249S cohorts. (B) Representative images and quantitation of tumor cryo-section stained with Ki67 in 4T1 PLVX and p53R249S tumors. (C) 5 x 10^4^ 4T1 cells expressing PLVX or p53R249S were injected onto the immunodeficient NOD/SCID mice and the image depicts similar tumor size between the two cohorts (n=4). Quantification graphs: FoV= Field of View, In all panels, error bars represent mean with standard error. In scatter dot plots, each dots represent one mice, p values are based on Student’s t test. ^∗∗^p < 0.01, ^∗^p < 0.05, ns=non-significant.

**Figure S7:**
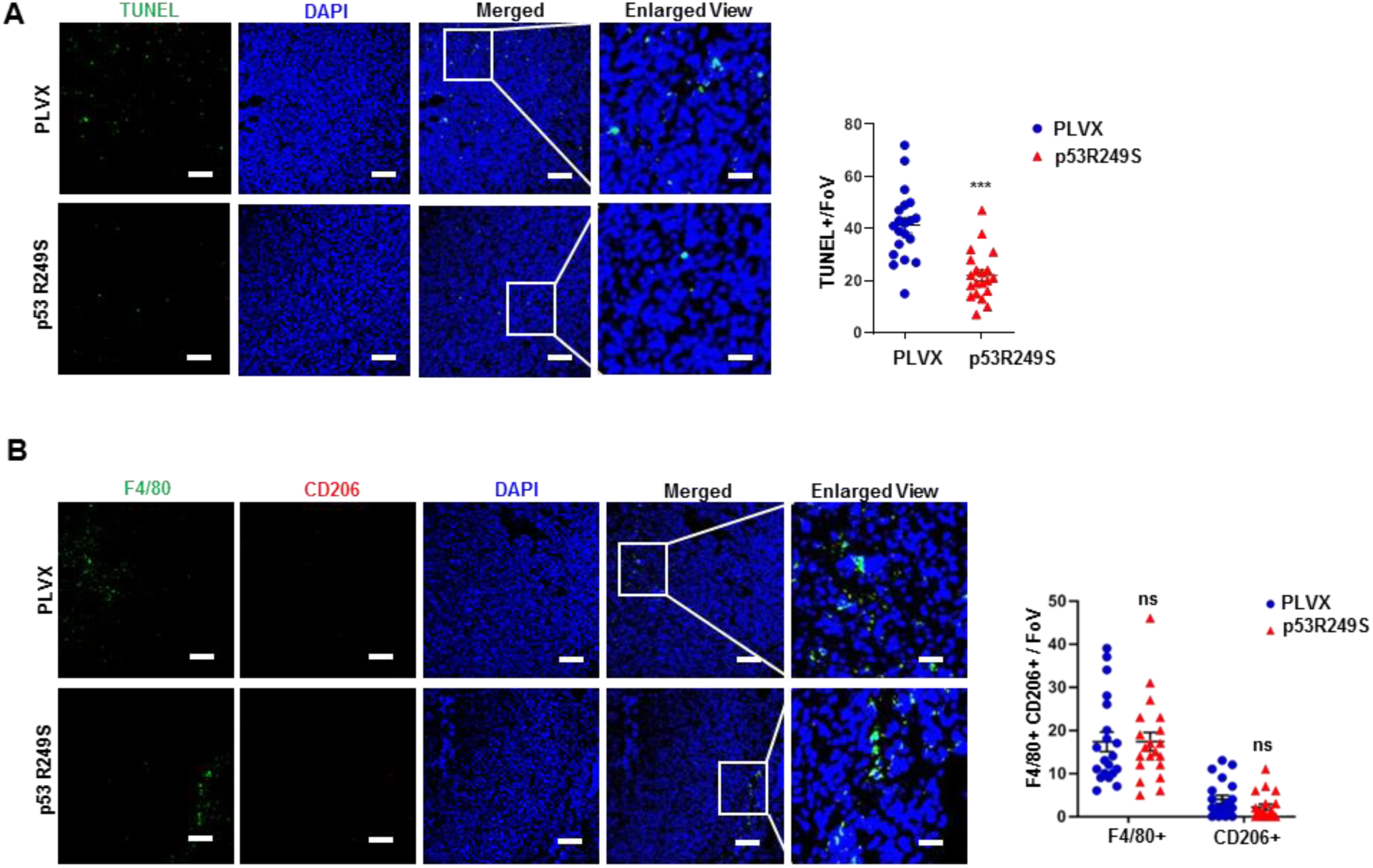
Mutant p53 suppress apoptosis in 4T1 tumor section. (A) Representative immunostaining for tumor cell death (TUNEL) in 4T1 PLVX and p53R249S tumor sections in Balb/C mice. (Right) Representative graph showed quantification of TUNEL positive cells. (B) Representative confocal images depicting the F4/80^+^CD206^+^ M2 type of TAMs in PLVX and p53R249S expressing tumors isolated from NOD/SCID mice on Day 21. (Right) Representative graphs indicate quantitation of F480+, CD206+ M2 macrophage subsets in NOD/SCID mice (right) (n=20). Images were captured at 20X, scale bar 25 μm except in enlarged panel which is 100 μm. Quantification graphs: In all panels, error bars represent mean with standard error. p values are based on Student’s t test. ***p < 0.001, ^∗^p < 0.05, ns=non-significant.

**Figure S8:**
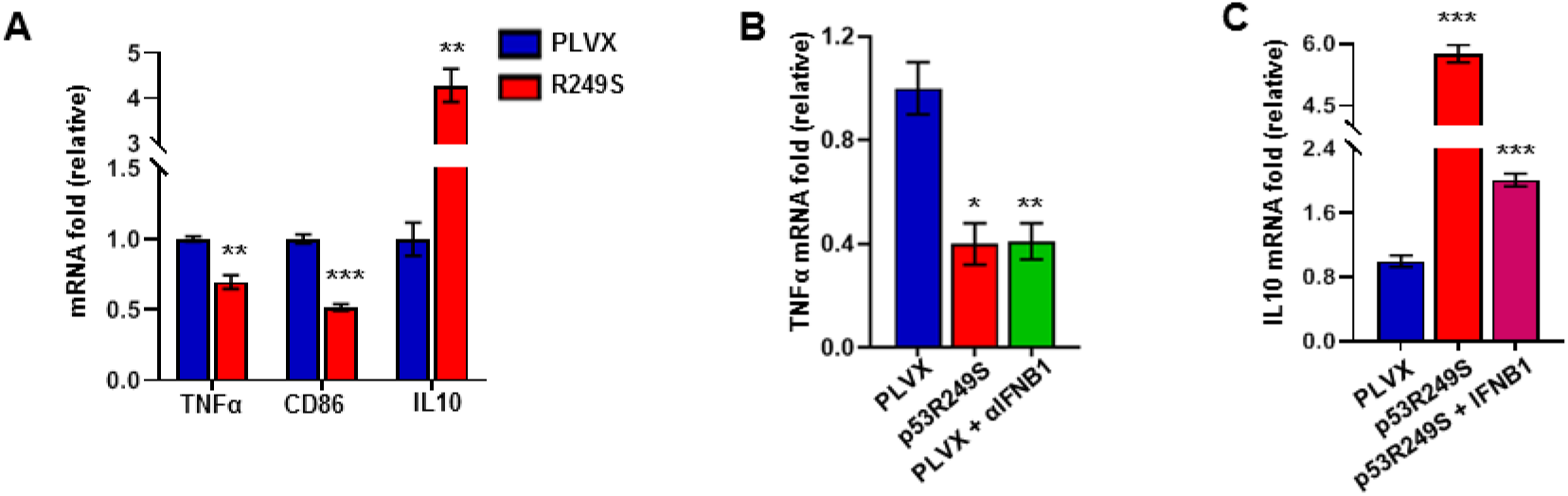
Mutant p53 alters RAW264.7 macrophage polarization *in vitro*. (A) Quantification of the expression of the indicated gene mRNA of RAW264.7 cells when cultured either in PLVX or p53R249S expressing 4T1 cell conditioned media. (B) RT-PCR analysis of TNFα mRNA expression in RAW264.7 when cultured in PLVX conditioned media containing anti-αIFNB1 antibody. (C) RT-PCR analysis of IL-10 expression in RAW264.7 when cultured in p53R249S conditioned media containing IFNB1. Quantification graphs: Error bars represent mean with standard deviation. p values are based on Student’s t test. ^∗∗∗^p < 0.001, ^∗∗^p < 0.01, ^∗^p < 0.05, ns=non-significant

**Figure S9:**
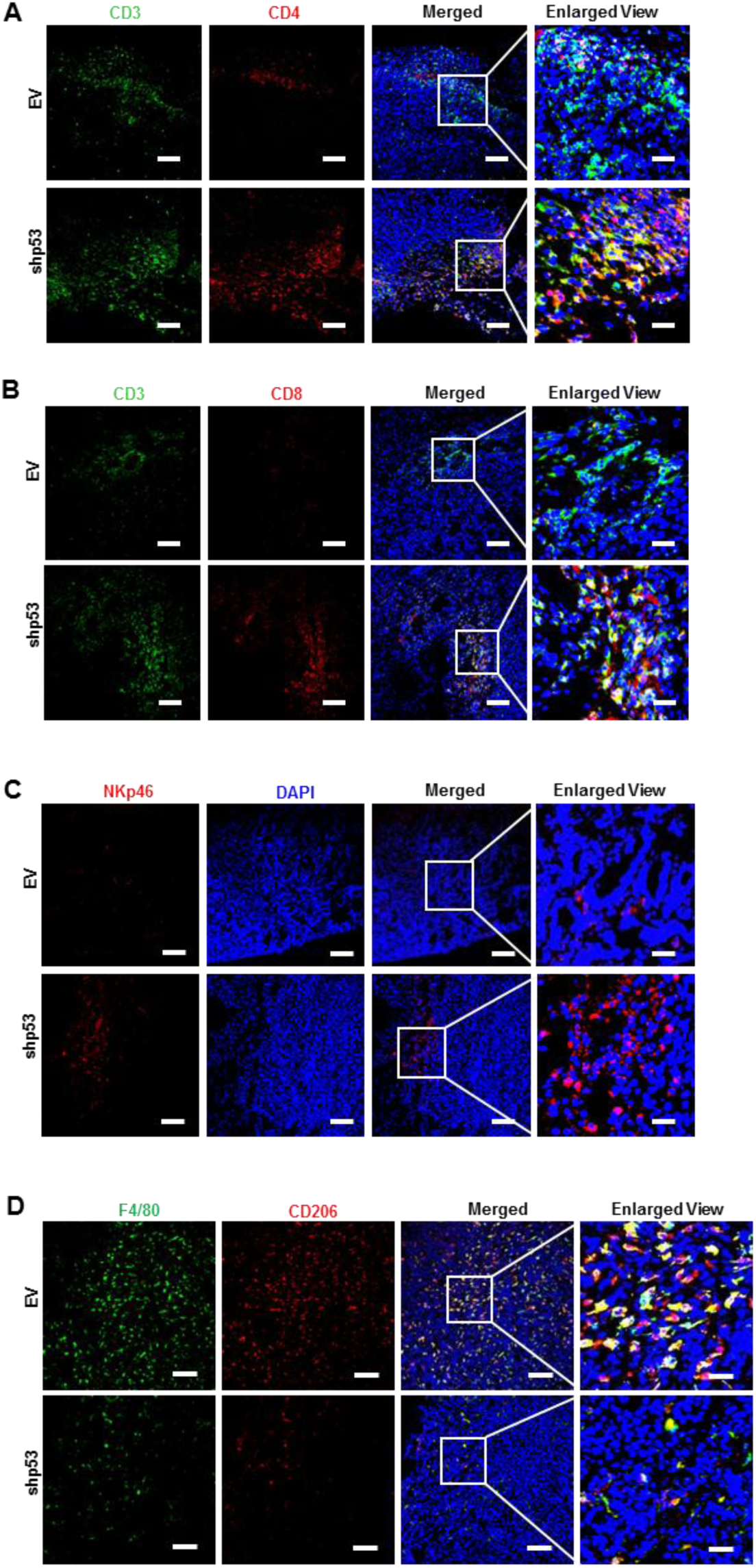
Mutant p53 knockdown promotes immune cells recruitment to restrict KPC tumor growth. Representative confocal micrographs of A) CD3^+^CD4^+^ T-helper and (B) CD3^+^CD8^+^ T-cytotoxic lymphocyte infiltration in KPC induced EV or shp53 tumor sections. (C) Representative confocal micrographs depicting the expression of NKp46 in EV or shp53 induced KPC tumors on Day 21. (D) Representative confocal images depicting the F4/80^+^CD206^+^ M2 type of TAMs in EV and shp53 tumors isolated from C57BL/6 mice on Day 21. All Images were captured at 20X, scale bar 25 μm except in enlarged panel which is 100 μm.

**Figure S10:**
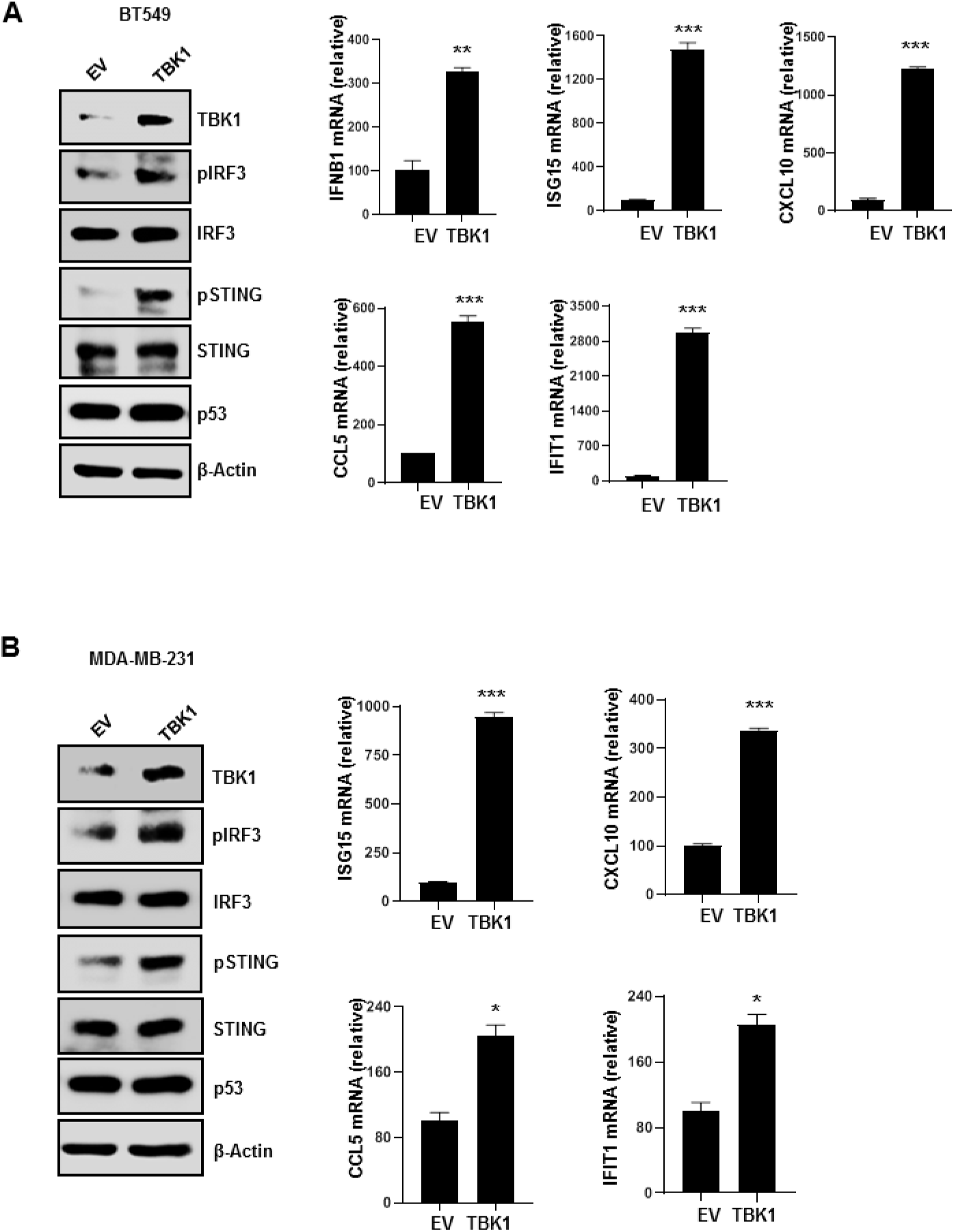
TBK1 overexpression activates DNA sensing pathway. (A) and (B) Immunoblots showing that overexpression of TBK1 in mutant p53 harboring BT549 and MDA-MB-231 cells leads to the phosphorylation of IRF3 and STING. Quantitative RT-PCR showing the increased expression of the indicated tumor suppressive cytokines upon TBK1 overexpression.

**Figure S11:**
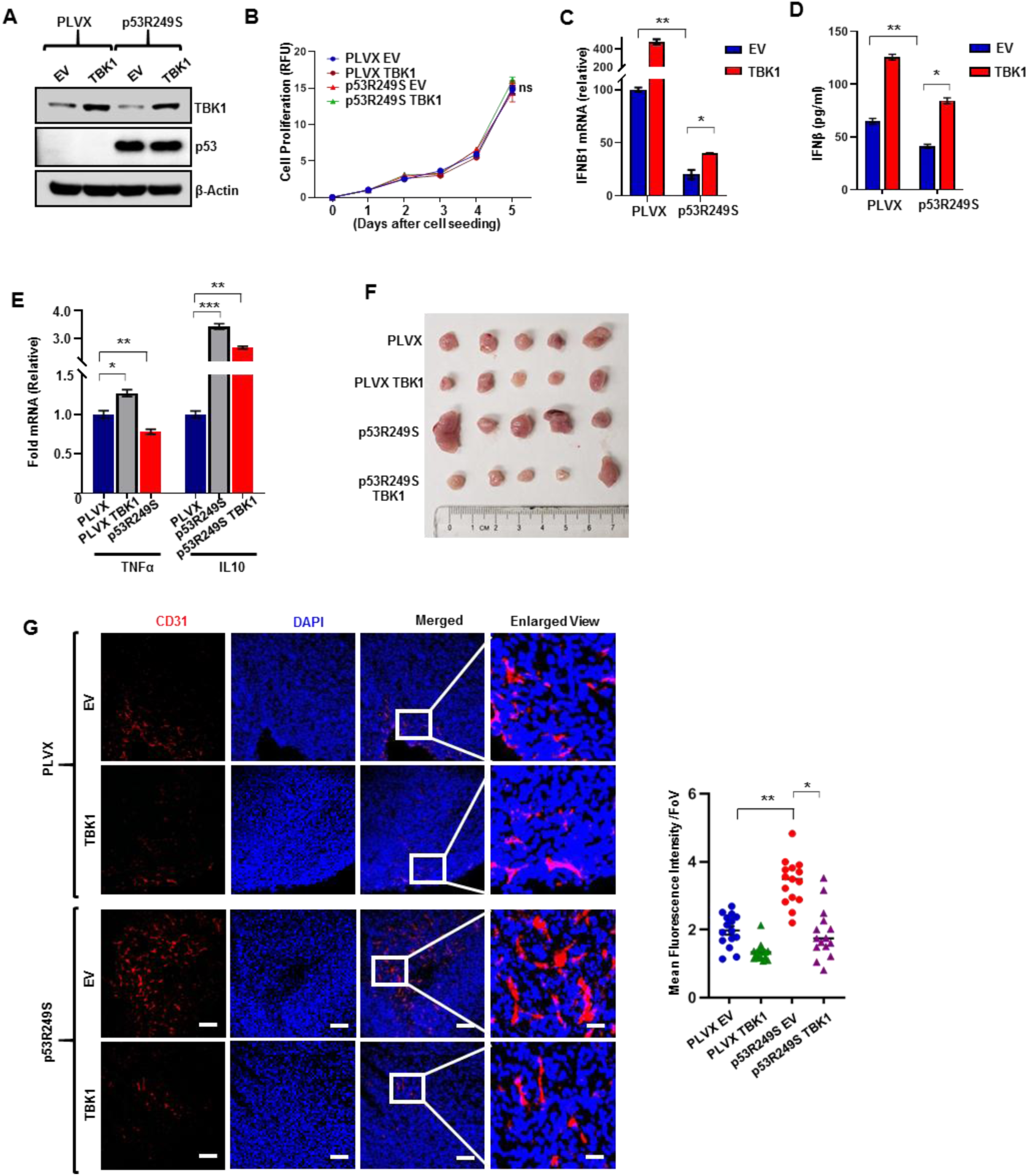
TBK1 overexpression inhibits tumor progression and downregulates neoangiogenesis. (A) 4T1 PLVX or p53R249S expressing 4T1 cells were induced with doxycycline for 24 hrs to induce TBK1. Immunoblots representing TBK1 expression status within the cells. (B) Representative graphical quantification of *in vitro* cell proliferation between 4T1 PLVX and p53R249S induced TBK1 cohorts. (C) Quantitative RT-PCR analysis of IFNB1 mRNA expression in the indicated cells. (D) Graph represents quantitative IFNB1 secretion as analyzed using ELISA. (E) Quantification of the expression of mRNA of indicated genes in RAW264.7 cells when cultured in PLVX and p53R249S induced TBK1 expressing 4T1 cell conditioned media. (F) 5 x 10^4^ 4T1 PLVX and p53R249S cells were injected into BALB/c mice and 20 mg/kg of Doxycycline was administered orally to induce TBK1 (n=5). Image depicts tumor volume of the indicated cohorts. (G) Confocal micrographs and graphical quantification depicting CD31 expression in the respective tumor cryo-sections (n=15). Graph at the right indicates quantification of CD31 intensity in the indicated cohorts. Quantification graphs: In Fig. B, C, D and E error bars represent mean with standard deviation. p values are based on Student’s t test. ^∗∗∗^p < 0.001, ^∗∗^p < 0.01, ^∗^p < 0.05, ns=non-significant. Images were captured at 20X, scale bar 25 μm except in enlarged panel which is 100 μm.

**Figure S12:**
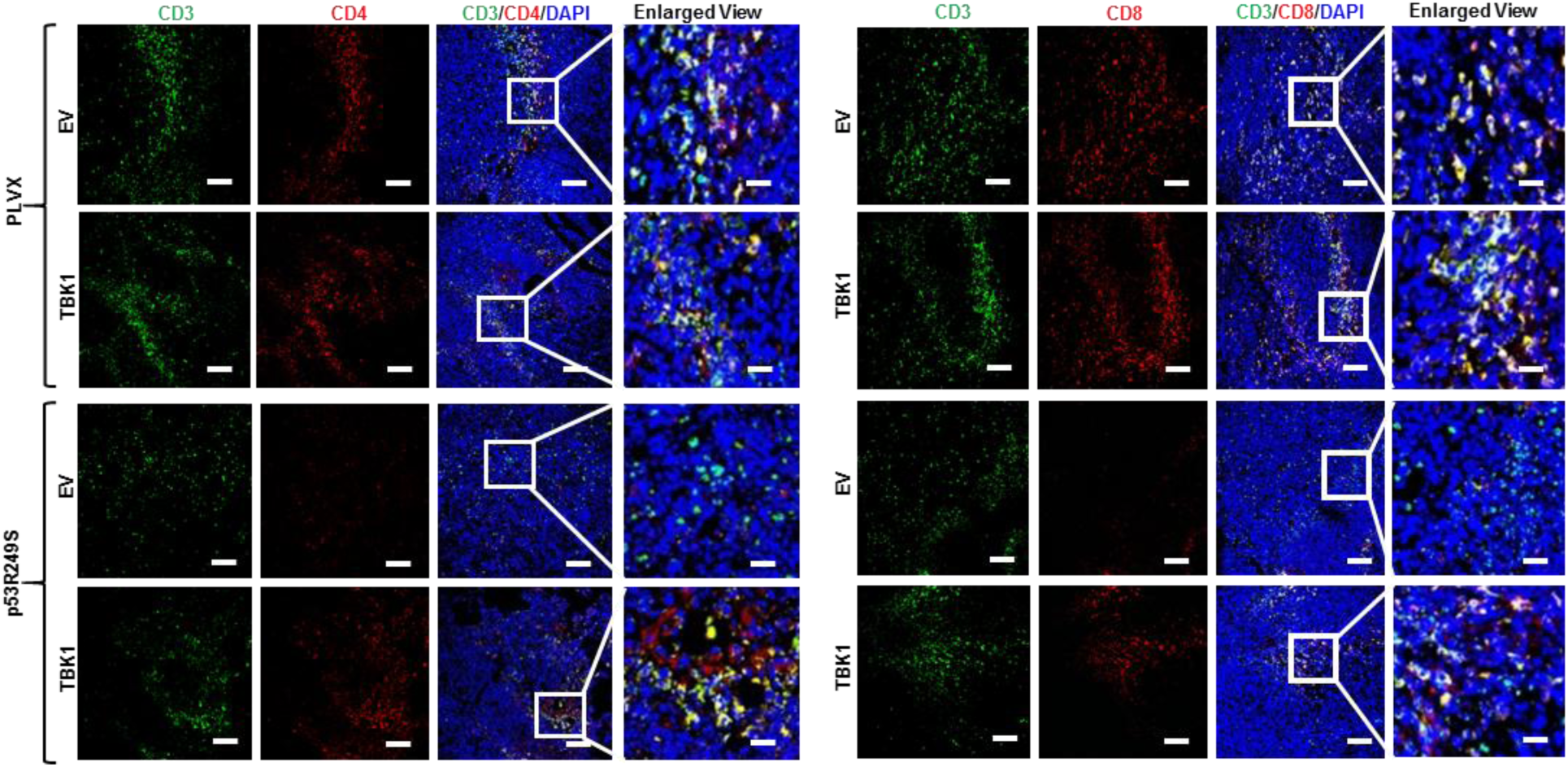
Representative confocal micrographs depicts recruitment of A) cytotoxic CD3^+^CD4^+^ T-helper cells and (B) Cytotoxic CD3^+^CD8^+^ T-cells infiltration to the tumor microenvironment in the indicated tumor sections. Images were captured at 20X, scale bar 25 μm except in enlarged panel which is 100 μm.

**Figure S13:**
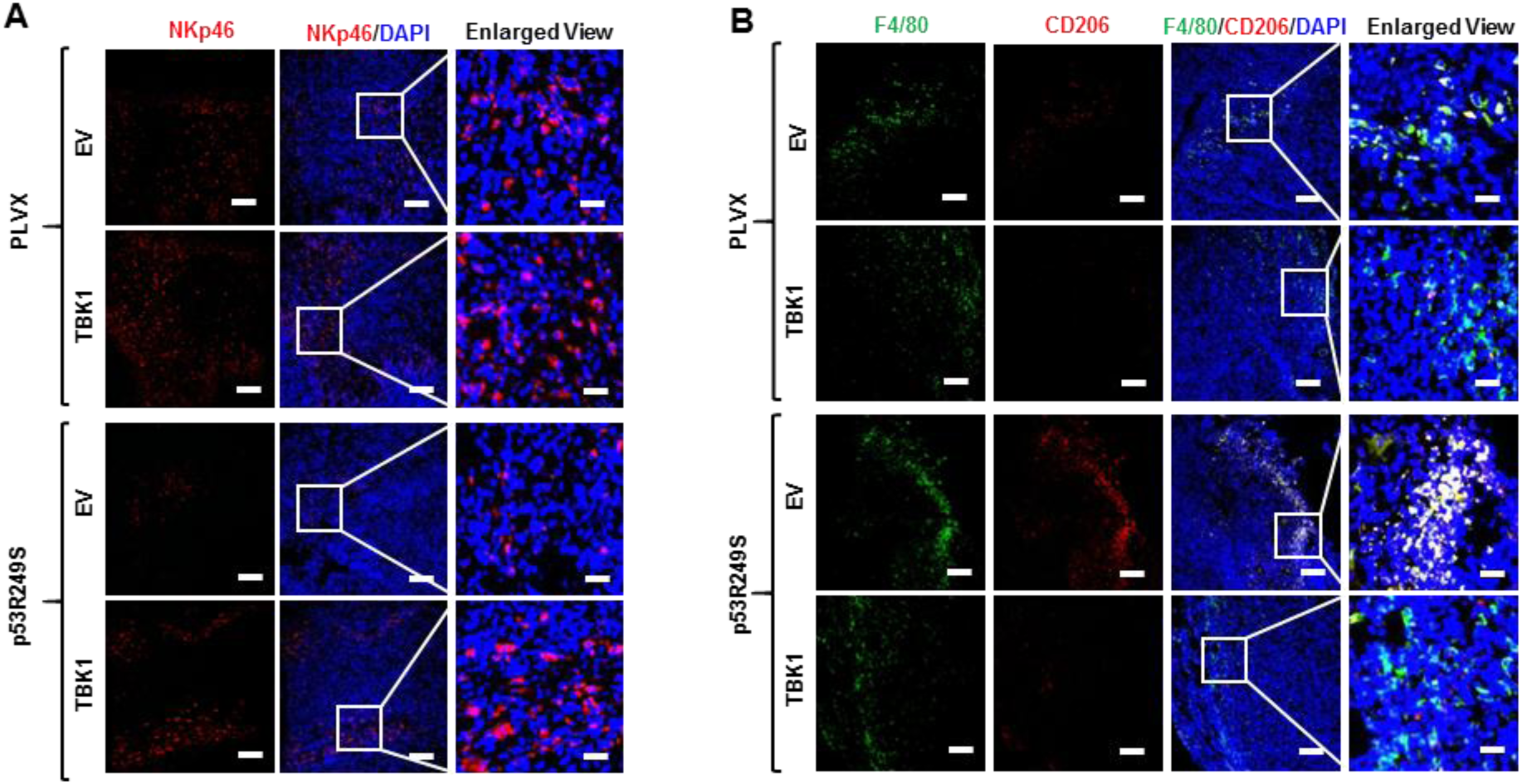
Representative confocal micrographs of A) NKp46^+^ NK-T-cells and (B) F480^+^CD206^+^ M2 macrophage infiltration in PLVX and p53R249S tumor sections.

