## Supplementary Materials for "Mutant p53 suppresses innate immune signaling to promote tumorigenesis"

**STAR Methods**

1. **Chemicals and Reagents.**

Lipofectamine 2000 was purchased form Invitrogen (Cat. 2082816). cGAMP (Cat. GA3-40-02), dsDNA (Cat. ECD-40-01) and poly(I:C) (31852-29-6) were purchased from Invivo Gen, HT-DNA (Cat. D6898), Doxycycline hyclate (Cat. D989were obtained from Sigma, LPS (cat. ALX-581-200-L001) was purchased from Enzo. Hoechst 33342 and DAPI were purchased from Life technology. Annexin-V FITC and PI were purchased from Biolegend. Doxycycline hydrochloride (Cat. BP2653-5) was purchased from Fisher Bioreagent. Halt protease and phosphatase inhibitor (Cat. TL274884) was obtained from Thermo. Sso Advanced universal SYBR Green super mix (172-5274) was purchased from Bio-Rad. Fluoromount G was purchased from SuthernBiotech and ProLong Gold Antifade Reagent was purchased from Invitrogen. Protein G agarose was obtained from KPL.

2. **Cloning and Plasmids**

PLKO and PLKO-shp53 were a kind gift from Robert Weinberg (Addgene #8453 and #19119). Non-targeting control and mouse p53 shRNA were cloned into EZ-Tet-pLKO-Puro (addgene:85966).

IRF3 knockout BT-549 and MD-MB231 cells lines were prepared using CRISPR/Cas9 technique. Guide RNA was cloned in lentiCRISPRv2-puro (addgene: 98290) as previously described (Shalem*, Sanjana*, et al., Science 2014). The viral particle was generated and infected to the target cells as described above and selected for 10 days.

pCMV-Myc-TBK1 was generated by PCR amplification of TBK1 from pWZL Neo Myr Flag TBK1 (Addgene #20648, gift from Jean Zhao) and cloning in frame with the MYC tag in PCMV-MYC (Clontech). pCMV-HA-STING was generated by PCR amplification of STING from IMR90 lung fibroblast cDNA and cloning into pCMV-HA (Clontech). pcDNA3-GFP-IRF3 was a kind gift from Nancy Reich (Stony Brook University). TBK1 was amplified from pCMV-MYC-TBK1 to clone into PLVX-puro (Clontech). The p53R249S cDNA was cloned into the lentivirus vector PLVX-puro (or hygro). p53 point mutants: Different p53 point mutants (HAp53P142L, HAp53P152Q, HAp53A161V, HAp53C174Y, HAp53R175H, HAp53R248W, HAp53R249S, HAp53R273H, HAp53R280K) were generated using site directed mutagenesis (NEB) of pCMV-HA-wildtype p53. A series of deletion mutants were prepared based on p53R249S, HAp53R249SΔ21-71, HAp53R249SΔ72-122, HAp53R249SΔ123-173, HAp53R249SΔ174-224, HAp53R249SΔ225-275, HAp53R249SΔ276-326, HAp53R249SΔ327-377 and cloned onto pCMV vector.

All the constructs used in the study were confirmed by DNA sequencing. The sequences for sgRNAs, shRNAs and primers for PCR mutagenesis used in this study are listed in Table 1.

3. **Cell culture.**

All the cell lines were purchased from ATCC and cultured according to the manufacturer’s instructions. MDA-MB-231, BT549, H1299, MiaPaCa2, A549 cells were cultured in complete Roswell Park Memorial Institute (RPMI) medium supplemented with 10% Fetal Bovine Serum (FBS). KPC (kind gift from Dr. Richard Linn, Stony Brook University), HEK293T, 4T1, RAW264.7 cells were cultured in complete Dulbecco's Modified Eagle Medium (DMEM) medium. MEFs (p53^-/-^, p53^R172H/R172H^) were isolated in Dr. Iwakuma's Lab, University of Kansas Medical Center, according to the protocol described earlier (A. Parrales et. al. *Nature Cell Biology*, 2016) and were cultured in complete DMEM medium. IMR-90 and HFF (kind gift from Dr. Lina Obeid and Dr. Jeffrey Stith lab, Stony Brook University) cells were cultured in DMEM with 15% FBS supplemented with non-essential amino acids.

4. **Lentiviral particles production in 293FT cells and stable gene knockdowns.**

For shRNA knockdown lentiviral particle was generated by transfecting 293T cells with 1.5 µg of ps-Pax2 (addgene), 0.5 µg of pCMV-VSV-G (addgene) and 2 µg of plasmid of interested genes using Lipofectamine 2000. Viral supernatant was collected post 48 hrs and 72 hrs of transfection. Target cells were infected with the viral particle using polybrene (5 µg/mL). Post 48 hrs after infection media was changed and selected with puromycin or hygromycine for 3 days.

5. **Generation of IRF3 knock out cells using CRISPR-Cas9**

To generate lentiviruses for transduction, HEK293T cells were transfected with plasmid(s) encoding IRF3 sgRNA and packaging vectors (VSVG and psPAX2) using a standard Lipofectamine 2000 transfection method. Lentiviruses encoding Cas9 were generated using the same technique. Culture supernatants were collected at 48 and 72 h post-transfection and used for infection of BT549 and MDA-MB-231 cells with polybrene (5 µg/ ml). Cells were selected with puromycin (1 µg/ml) 48 h post infection for 10 days.

6. **Cell proliferation assay.**

4T1 PLVX or p53R249S cells (3,000) and 4T1 PLVX or p53R249S induced TBK1 cells were seeded on a 96-well plate and cell proliferation was detected for next five days. Viable cells were measured by CellTiter-Blue® Cell Viability Assay kit (Promega) according to the manufacturer protocol. Briefly, 20ul of cell titre blue reagent was directly added to the culture medium and incubated at 37^0^C for 4 h and plates were shaken for 10 sec and the fluorescence reading were obtained by reading the plate at 570/590 nm by Molecular Device Spectra Max M5 instrument.

7. **RNA isolation, cDNA synthesis, and real-time quantitative PCR (qRT-PCR).**

Total RNA from cells/tissues were collected in RLT buffer and was isolated using the Qiagen mini RNA isolation kit. RNA quantity and quality were confirmed with a NanoDrop ND-1000 spectrophotometer, cDNA was synthesized using 500ng of total RNA using oligo (dT) primers and Reverse Transcriptase (Quanta). Real-time qRT-PCR was performed in Bio-Rad CFX96 Touch real-time PCR detection system using Universal SYBr Green Supermix (Bio-Rad). Gene-specific primers sequences listed in Supplementary Table 1.

8. **IFN Beta measurement by ELISA.**

p53KD BT549, KPC and p53^-/-^, p53 ^R175H/ R175H^ MEFs and p53R248W overexpressing H1299 and p53R249S expressing 4T1 cells (10^6^) were seeded on a 60mm dish and after 24 hrs cells stimulated by HT-DNA (4 µg) for another 18hrs and collected supernatants were analyzed using VeriKine Human IFN Beta (pbl Assay Science 41410-1) or mouse IFN Beta ELISA Kit (pbl Assay Science 42400-2). Quantification of IFNB1 concentration was performed in triplicates according to the manufacturer protocol and the reading was taken at 450 nm by Molecular Device Spectra Max M5 instrument and calculated using an IFNB1 standard curve.

9. **Immunofluorescence microscopy.**

GFP-IRF3 positive H1299 or MDA-MB-231 cells were grown onto 1% gelatin pre-coated glass coverslips and after all the treatment cells were washed twice with DPBS and counter stained with Hoechst 33342 and mounted with the ProLong Gold Antifade Reagent. Images were captured with a Nikon T*i* epifluorescence microscope and processed using Nikon AR software.

10. **Flow cytometry.**

P53KD, IRF3KO or Double KD BT549 cells and p53R248W overexpressing H1299 cells were transfected with HT-DNA for 24hrs and induced apoptosis was determined using flow cytometric analysis. Cells were washed with PBS, resuspended in 100 μl of binding buffer and further incubated with Annexin-V FITC and PI for 15 min in dark at room temperature. Prior to flow cytometric analysis, 400 μl of binding buffer was added and immediately subjected for the FACS analysis for the number of apoptotic cells. Data was generated using BD FACS Calibure and analyzed using BD FACS DIVA 6.2 software.

11. **Immunoblotting and Immunoprecipitation.**

To prepare cell lysates for western blotting, the cells were lysed on the dish using RIPA (0.5% SDS, 0.1% Sodium Deoxycholate, 0.5% NP40, 1 mM EDTA, in PBS pH 7.4 and filter-sterilize) buffer supplemented with protease and phosphatase inhibitors, scraped and placed into microcentrifuge tubes, sonicated and centrifuged at 13,000g for 10 mins at 4 °C to remove insoluble material. Protein concentration was determined using the Micro BCA Protein Assay kit (Pearce) and equal amounts of protein were resolved on 8 or 10% Bis-Tris polyacrylamide gels, transferred to a PVDF membrane blocked with 5% milk and incubated with primary antibody over night at 4^0^C. For co-immunoprecipitation of proteins, cells were washed with PBS, harvested and lysed in immunoprecipitation buffer (50 mM Tris-HCl pH 8.0, 150 mM NaCl, 0.05 mM EDTA, 1% NP40 and 10% glycerol). Lysate was clarified by centrifugation at 20,000g (4 °C) for 20 min, pre-cleared with protein-G agarose (KPL) for 2 h at 4 °C and then immunoprecipitated overnight with the corresponding antibodies. For cell fractionation assay, mutant p53 knockdown KPC and BT549 cells were lysed, cytosolic and nuclear fractions were extracted using NE-PER™ Nuclear and Cytoplasmic Extraction kit (Thermo Scientific, 78835) according to the manufacturers protocol. All antibody information can be found in Supplementary Table 2.

12. **Macrophage polarization Assay.**

RAW 264.7 mouse macrophages were seeded in a 6-well plate (10^5^) after 24 hrs cultured medium was replaced with 4T1 PLVX or p53R249S conditioned medium containing either control antibody/αIFNB1 antibody or IFNB1 protein for another 24 hrs. After all the incubation cells were harvested, RNA was isolated and further processed for RT-PCR analysis.

13. ***In vivo* animal experiments.**

4-6 weeks old female BALB/c, NOD/SCID mice were purchased from Charles River. Mice were anesthetized using Isoflurane and 4T1 PLVX or p53R249S cells (50,000) in 0.1 ml PBS were injected in the mammary gland after anaesthetizing the mice. Mice were monitored and tumor volume was measured manually using slide calipers every other day till day 21 when all the mice were sacked.

4-6 weeks old male c57BL6 mice were purchased from Jackson Laboratory. KPC inducible EV or shp53 cells were trypsinized washed twice with PBS and 1X10^5^ cells in PBS were injected subcutaneously at the back after anaesthetizing the mice. All the mice were given doxycycline (20 mg/kg) orally every other day starting from day 4 to induced either EV or shp53. Tumors were monitored and volume was measured manually using slide calipers till day 21 when all the mice were sacked and the tumors were processed for further experiment.

For the TBK1 rescue experiment, 4-6 weeks old female BALB/c mice were used. 4T1 PLVX (Inducible EV or TBK1) and p53R249S (Inducible EV or TBK1) cells were trypsinized, washed twice with PBS and 50,000 cells in PBS were injected in the mammary gland after anaesthetizing the mice. Mice were monitored and were given orally 20 mg/kg Doxycycline every other day to induce either EV or TBK1. Tumor volume were measured manually using slide calipers till day 21 when all the mice were sacked and the tumors were resected and further processed.

All the experiments with mice were conducted in Stony Brook University animal care facility and in accordance with the Institutional Animal Care and Use Committee.

14. **Immunohistochemistry.**

For the immunohistochemistry assays, the tumors were resected and fixed in 4% paraformaldehyde for overnight, dehydrated with a gradient sucrose solution of 15% and then 30% at RT. Then tumor tissues were embedded in optimal cutting temperature compound (OCT) and immediately frozen at −80°C until further use. Tumor blocks were cryo-sectioned at a thickness of 10μm using Leica Cryostat (Leica CM1900). Tumor sections were washed thrice with PBS to wash residual OCT compound, permeablized with 0.1% Triton X-100, blocked in 1% BSA in PBS for 45 mins at RT and incubated with the primary antibodies against: anti-Ki67, anti-CD3, anti-CD4, anti-CD8, anti-F4/80, anti-CD206, anti-NKp46 and anti-CD31 over night at 4^0^C. After the primary incubation sections were washed and incubated with Alexa flour 555 or Cy5-labelled secondary antibodies for one hour at room temperature, tumor sections were washed three times and counter stained with DAPI and mount with Fluoromount G. Immunohistochemical images of Ki67 were captured in Nikon Ti microscope and immunofluorescence images were captured using Leica TCS SPF5 II confocal microscope at 20X magnification and analyzed using ImageJ software. A list of all antibodies can be found in Table 2.

15. **TUNEL Staining**.

TUNEL analysis was performed in 4T1 PLVX and p53R249S tumor sections using the Apoptag Fluorescein In Situ Apoptosis Detection Kit (Sigma Millipore; S7110) according to manufacturer’s instructions. Briefly, tissue cryo-sections were washed in DPBS thrice to remove residual OCT medium, sections were then covered with equilibration buffer for a minimum of 1 min. Then the sections were incubation at 37°C for 1 hr with TdT enzyme (30% enzyme and 70% reaction buffer), followed by 10 mins incubation at room temperature in stop/wash buffer. Sections were then incubated in fluorescein anti-digoxigenin conjugated secondary antibody, washed, counter stained with DAPI and mounted.

16. **Statistical analysis.**

The statistical differences in all assays including Fold difference in mRNA, cell proliferation and growth, flow cytometry and tumor growth between different samples and/or treatments were analyzed by two-tailed Student’s t-tests using Microsoft Excel 2007 and all the graphs were made on GraphPad Prism 8 (GraphPad Software). Statistical significance was set at P < 0.05, unless otherwise stated in the text. All experiments were carried out with at least three biological replicates otherwise mentioned in the figure legend. The numbers of animals used are described in the corresponding figure legends.

**Table S1:** List of Primers

| Gene name | Forward Primer | Reverse Primer | Comments |
| --- | --- | --- | --- |
| IRF3 #2 | CACCGCCAGTGGTGCCTACACCCCG | AAACCGGGGTGTAGGCACCACTGGC | sgRNA targeting human IRF3 |
| IRF3 #3 | CACCGGCAACACTTCTTTCCGGTTC | AAACGAACCGGAAAGAAGTGTTGCC | sgRNA targeting human IRF3 |
| p53 (m) | CTAGCGTACATGTGTAATAGCTCCTACTAGTGGAGCTATTACACATGTACTTTTTG | AATTCAAAAAGTACATGTGTAATAGCTCCACTAGTAGGAGCTATTACACATGTACG | shRNA targeting mouse p53 |
| PLVXTBK1 | ATATCTCGAGATGCAGAGCACTTCTAATCATCTG | ATATGCGGCCGCCTAAAGACAGTCAACGTTGCGA | Sub cloned in PLVX |
| MYC-TBK1 | ATATGTCGACCATGCAGAGCACTTCTAATCATCTG | ATATGCGGCCGCCTAAAGACAGTCAACGTTGCGA | Sub cloned in PCMV |
| HA-STING | ATATGCGGCCGCCTAAAGACAGTCAACGTTGCGA | ATATGCGGCCGCTCAAGAGAAATCCGTGCGG | Sub cloned in PCMV |
| HA-p53R249SΔ21-71 | CCCGTGGCCCCTGCACCA | TGAAAATGTTTCCTGACTCAGAGGGGGCTC | Site directed mutagenesis |
| HA-p53R249SΔ72-122 | ACTTGCACGTACTCCCCTG | GGGAGCAGCCTCTGGCAT | Site directed mutagenesis |
| HA-p53R249SΔ123-173 | AGGCGCTGCCCCCACCAT | CACAGACTTGGCTGTCCCAGAATGC | Site directed mutagenesis |
| HA-p53R249SΔ174-224 | GTTGGCTCTGACTGTACC | CACAACCTCCGTCATGTG | Site directed mutagenesis |
| HA-p53R249SΔ225-275 | GCCTGTCCTGGGAGAGAC | CTCAGGCGGCTCATAGGG | Site directed mutagenesis |
| HA-p53R249SΔ276-326 | TATTTCACCCTTCAGATCC | ACAAACACGCACCTCAAAG | Site directed mutagenesis |
| HA-p53R249SΔ327-377 | TCCCGCCATAAAAAACTC | TTCTCCATCCAGTGGTTTC | Site directed mutagenesis |
| IFNβ (h) | GTCAGAGTGGAAATCCTAAG | TATGCAGTACATTAGCCATC | qPCR primers |
| IFIT1 (h) | TACAGCAACCATGAGTACAA | TCACATAGGCTAGTAGGTTG | qPCR primers |
| CCL5 (h) | AGCAGTCGTCTTTGTCAC | TAGCTCATCTCCAAAGAGTT | qPCR primers |
| CXCL10 (h) | TACCTGCATCAGCATTAGTA | TGTAGCAATGATCTCAACAC | qPCR primers |
| ISG15 (h) | GAACTCATCTTTGCCAGTA | ATCTTCTGGGTGATCTGC | qPCR primers |
| IL6 (h) | CCCCCAATAAATATAGGACT | GATAGAGCTTCTCTTTCGTT | qPCR primers |
| IFNβ (m) | AAGATCAACCTCACCTACAG | AAAGGCAGTGTAACTCTTCT | qPCR primers |
| CCL5 (m) | AGTGGGTTCAAGAATACATC | CTAGGACTAGAGCAAGCAAT | qPCR primers |
| CXCL10 (m) | AAGTTTACCTGAGCTCTTTT | AGTATCTTGATAACCCCTTG | qPCR primers |
| ISG15 (m) | ACAGTGATGCTAGTGGTACA | AAGACCTCATAGATGTTGCT | qPCR primers |
| IL6 (m) | GTCTTCTGGAGTACCATAGC | TATCTGTTAGGAGAGCATTG | qPCR primers |
| TNFα | ATCTTCTCAAAATTCGAGTG | ACCACTAGTTGGTTGTCTTT | qPCR primers |
| CD86 | GAACAACCAGACTCCTGTAG | GTCACAAAGATAAGGATTGC | qPCR primers |
| IL10 | TTACCTGGTAGAAGTGATGC | TGTAGACACCTTGGTCTTG | qPCR primers |

| **Sl. No.** | **Antibody** | **Company** | **Cat. No.** | **Species** | **Clone** |
| --- | --- | --- | --- | --- | --- |
| 1. | Anti-STING | Cell Signaling | 136475 | Rabbit | monoclonal |
| 2. | Anti-TBK1 | Cell Signaling | 35045 | Rabbit | monoclonal |
| 3. | Anti-IRF3 | Cell Signaling | 4302 | Rabbit | monoclonal |
| 4. | Anti-pSTING | Cell Signaling | 197815 | Rabbit | monoclonal |
| 5. | Anti-pTBK1 | Cell Signaling | 5483T | Rabbit | monoclonal |
| 6. | Anti-pIRF3 | Cell Signaling | 29047S | Rabbit | monoclonal |
| 7. | Anti-P53 | Santa Cruz | Sc126 | Mouse | monoclonal |
| 8. | Anti-P53 | Santa Cruz | Sc6243 | Rabbit | monoclonal |
| 9. | Anti-GAPDH | Genetex | GTX100118 | Rabbit | polyclonal |
| 10. | Anti-β Actin | Sigma | A3854 | Mouse | monoclonal |
| 11. | Anti-Myc | Santa Cruz | Sc40 | mouse | monoclonal |
| 12. | Anti-HA | Santa Cruz | Sc7392 | mouse | monoclonal |
| 13. | Anti-GFP | Santa Cruz | Sc9996 | mouse | monoclonal |
| 14. | Anti-Ki67 | Abcam | Ab15580 | Rabbit | polyclonal |
| 15. | Anti-NKp46 | Biolegend | 137607 | Rat | monoclonal |
| 16. | Anti-F4/80 | Biolegend | 123119 | Rat | monoclonal |
| 17. | Anti-CD3 | Invitrogen | 1989145 | Rat | monoclonal |
| 18. | Anti-CD4 | Invitrogen | 4348408 | Rat | monoclonal |
| 19. | Anti-CD8 | Invitrogen | 2089510 | Rat | monoclonal |
| 20. | Anti-CD206 | R & D systems | AF2535 | Goat | Polyclonal |
| 21. | Anti-CD31 | BD | 557355 | Rat | monoclonal |
| 22. | Anti-secondary Alexaflour555 | Invitrogen | 2026158 | Donkey | Polyclonal |
| 23. | Anti-secondary-Cy5 | Abcam | Ab6565 | Goat | polyclonal |
| 24. | Anti-Lamine B1 | Abcam | ab16048 | Rabbit | polyclonal |

**Table S2:** List of antibodies
